# Identification of a Lipid Oxygen Radical Defense pathway and its epigenetic control

**DOI:** 10.1101/2025.02.25.640160

**Authors:** Francisco S. Mesquita, Laurence Abrami, Romain Forey, Béatrice Kunz, Charlène Raclot, Lucie Bracq, Filipe Martins, Danica Milovanovic, Evarist Planet, Olga Rosspopoff, Didier Trono, F. Gisou van der Goot

## Abstract

Membrane phospholipids are vulnerable to oxidative radicals, and uncontrolled lipid peroxidation affects cell viability. Cells have evolved quality control and defense mechanisms, of which the genetic regulation is not fully understood. Here, we identify what we have coined the Lipid Oxygen Radical Defense (LORD) pathway. It is epigenetically repressed by a complex comprising the KRAB-zinc finger protein ZNF354A, the scaffold protein KAP1/TRIM28, the histone methyltransferase SETDB1, and the transcriptional activator ATF2. Upon lipid peroxide accumulation, p38– and JNK-dependent phosphorylation of ATF2, KAP1/TRIM28, and ZNF354A leads to disassembly of the repressive complex, releasing ZNF354A from specific DNA loci and activating a network of protective genes, including NRF2 targets. The pathway affects the cellular sensitivity to oxidative stress and ferroptosis, revealing a novel layer of epigenetic control in lipid quality control and damage repair. This positions the LORD pathway as a promising therapeutic target for diseases linked to chronic inflammation, neurodegeneration and cancer.

## INTRODUCTION

Reactive oxygen species (ROS) are inevitable byproducts of aerobic metabolism ^1,2^. These molecules play a dual role in cellular physiology: they are vital for signaling pathways ^1,2^, but excessive ROS accumulation leads to oxidative stress and cellular damage ^1–3^. Elevated hydrogen peroxide (H₂O₂) levels can trigger Fenton reactions with transition metals, such as ferrous iron (Fe²⁺), producing highly reactive hydroxyl radicals that inflict severe oxidative damage to biomolecules such as DNA, proteins, and lipids ^1,2^, particularly phospholipids containing polyunsaturated fatty acids (PUFAs) present in cellular membranes ^4–6^. Accumulation of lipid peroxides and their deleterious by-products is a hallmark of oxytosis or ferroptosis, a non-apoptotic form of cell death increasingly recognized for its importance in aging, cancer, cardiovascular disease, neurodegeneration, and chronic inflammation ^4,5,7–11^. Cells have evolved sophisticated antioxidant defense systems to protect against membrane lipid peroxidation, both selenium/thiol-dependent and independent pathways ^7,9,12,13^. How these quality control mechanisms are regulated at a transcriptional level, particularly in complex organisms like mammals ^14^, is not fully understood, and is the focus of the present study.

A complex network of transcription factors coordinates oxidative stress responses, with NRF2 (NF-E2-related factor 2) as the principal regulator ^2,9,14–16^. Controlled by the redox-sensitive cytoplasmic inhibitor Keap1, NRF2 binds to antioxidant response elements (ARE) in the 5′ UTR, driving the expression of antioxidant genes, xenobiotic transporters, and iron homeostasis regulators ^15–18^. NRF2 controls various detoxification systems ^7,15,17,19^, including the thiol-independent oxidoreductase FSP1 (ferroptosis suppressor 1)^20–22^ and factors required for glutathione (GSH) synthesis, such as SLC7A11, the cystine/glutamate antiporter crucial for cystine uptake ^23,24^. GSH is essential for the activity of GPX4 (glutathione peroxidase-4), the primary enzyme that reduces lipid hydroperoxides^13,25^. GPX4 is regulated by NRF1, highlighting functional overlap among transcription factors while maintaining distinct cellular targets ^26^. Other factors like NF-κB, HIF-1, FOXO1, p53, YAP/TAZ, and ATF4 are also involved in maintaining redox balance, influencing lipid peroxidation, inflammation, and hypoxia responses, ultimately affecting cellular sensitivity to oxidative stress ^2,14,27–31^. Redox-driven epigenetic modifications can further shape the expression of specific antioxidant genes^32–34^. However, despite this wealth of information, major gaps in understanding the transcriptional regulation of lipid quality control remain. The existence of a master regulator remains uncertain, and the mechanisms controlling specific antioxidant gene programs are yet to be elucidated.

In this study, we uncovered a defense pathway against ROS-triggered lipid damage, which we have termed the LORD pathway, for Lipid Oxygen Radical Defense. This pathway is under epigenetic regulation. DNA recognition is mediated by ZNF354A, a member of the largest family of mammalian transcriptional modulators, the Krüppel-associated box (KRAB) domain zinc finger proteins (KZFP) ^35,36^. We found that ZNF354A is part of a repressive complex containing the global repressor KAP1/TRIM28 (hereafter referred to as KAP1), the histone methyltransferase SETDB1, but also the transcriptional activator ATF2, which plays a crucial regulatory role. Under homeostatic conditions, the ZNF354A-containing complex represses transcriptional programs involved in lipid quality control, antioxidant defenses, and other response pathways. Accumulation of lipid peroxides leads to phosphorylation-dependent disassembly of the repressive complex, and release of ZNF354A from specific DNA loci, allowing activation of the antioxidant response pathway. This work highlights the underexplored dynamics of epigenetic repressive complexes and the genetic response to oxidative lipid damage. It also paves the way towards the identification of potential targets for therapeutic intervention in a variety of conditions such as chronic inflammation, infections, cancer, and aging-related disorders.

## RESULTS

### ZDHHC20L as a readout for the stress response pathway

In previous work on the S-acylation of the SARS-CoV-2 Spike protein by the acyltransferase ZDHHC20^37,38^, we found that infection induces a transcriptional shift in the *ZDHHC20* gene, generating N-terminally extended isoforms of the protein, which we here collectively term ZDHHC20 Long (ZDHHC20L). These longer isoforms are also expressed in cells exposed to the pore-forming toxin aerolysin and *in vivo*, upon chemical-induced colitis^37^. We additionally documented the presence of ZDHHC20L in untreated hepatocellular carcinoma cell line HepG2^37^, a finding we extended here to MCF7 breast cancer and HCT116 human colorectal carcinoma cells (Fig. 1a). We also investigated whether ZDHHC20L expression could be induced by aging, a process where increased stress pathways are linked to the gradual decline of repair mechanisms^39^. While in total cell extracts (TCEs) from primary lung epithelial cells of young donors only the canonical form of ZDHHC20 was detected, ZDHHC20L became increasingly expressed in cells from donors above 60 (Fig. 1b). ZDHHC20L expression is thus induced under a variety of conditions, ranging from some cancers to aging, viral infection, and other stresses. This study aimed to identify the stress response pathway that leads to ZDHHC20L expression.

**Figure 1.**
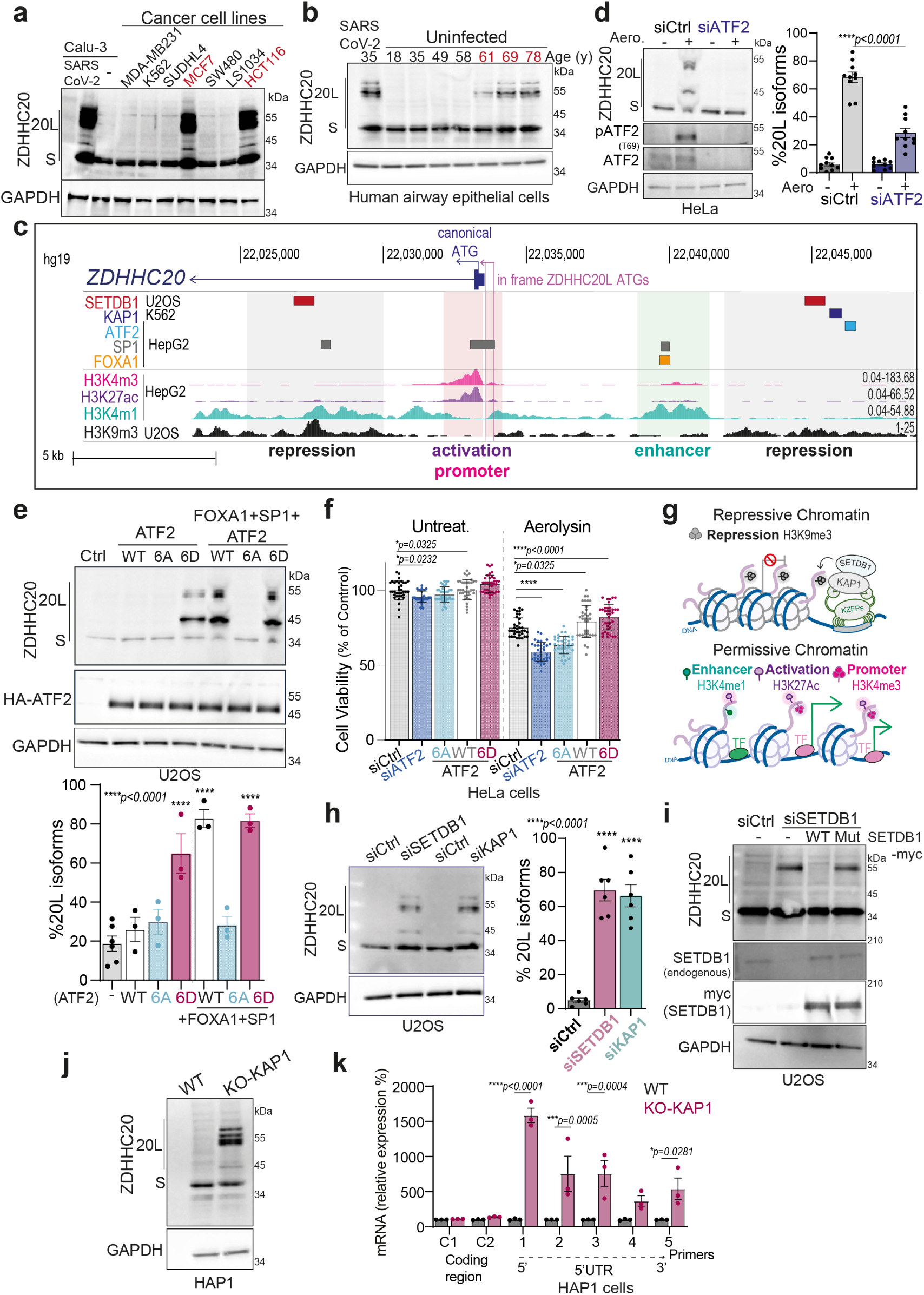
ATF2 and SETDB1-KAP1 dependent induction of ZDHHC20L expression. **a, b**, Western blot (WB) of ZDHHC20 isoforms (Short-S, 42 kDa; 20L, ∼49, 62, 69 kDa) and GAPDH (loading control) in a, indicated cancer cell lines and SARS-CoV-2-infected Calu-3 cells (MOI 0.1, 24 h); b, primary human airway epithelia from donors of different ages, untreated or SARS-CoV-2-infected (MOI 0.1, 48 h). **c,** Overview of the ZDHHC20 locus (first intron, exon, 5′ UTR; hg19). Tracks show in-frame start codons (ATGs), transcript (GENCODE), histone modifications (ENCODE: H3K9me3, repression; H3K4me1, enhancer; H3K27Ac, activation; H3K4me3, promoter), and SETDB1/KAP1, ATF2, SP1, FOXA1 ChIP-seq peaks. **d,** WB as in a, total and phospho-ATF2 in HeLa cells (siCtrl or siATF2), untreated or treated with proaerolysin (10 ng/ml, 1 h at 37 °C, incubated further 8 h). Quantification of 20L isoforms shown as mean ± SEM, dots represent independent experiments (n = 10); *p* values by two-way ANOVA, Sidak’s correction. **e,** WB as in a, in U2OS expressing ATF2 (WT, 6D, 6A), or FOXA1 and SP1. Results mean ± SEM, independent experiments (Ctrl, n = 6; others, n = 3); *p* values versus Ctrl, two-way ANOVA, Dunnett’s correction. **f,** Cell viability of HeLa cells (ATP assay), normalized to untreated siCtrl (100%), mean ± SD from one of three representative assays (n = 34 replicates); *p* values vs. siCtrl, two-way ANOVA, Dunnett’s correction. **g,** Schematic: KAP1-KZFPs recruit SETDB1, promoting H3K9me3-mediated repression and active histone mark removal (adapted from ref. 36). **h**, WB as in d, U2OS cells siCtrl, siSETDB1, or siKAP1-depleted (72 h). Results mean ± SEM, independent experiments (n = 6); *p* values vs. Ctrl, two-way ANOVA, Dunnett’s correction. **i,** WB as in h, siSETDB1 cells expressing WT or catalytically inactive SETDB1 mutant (Mut). **j,** WB as in a, in HAP-1 wild-type or KAP1-knockout cells. **k,** mRNA quantification across ZDHHC20 transcripts (coding regions: C1, C2; 5’UTR positions: 1–5) in HAP-1 wild-type or KAP1-knockout cells, mean ± SEM, independent experiments, *p* values vs. normalized WT, two-way ANOVA, Dunnett’s correction.

**Supplementary Figure 1.**
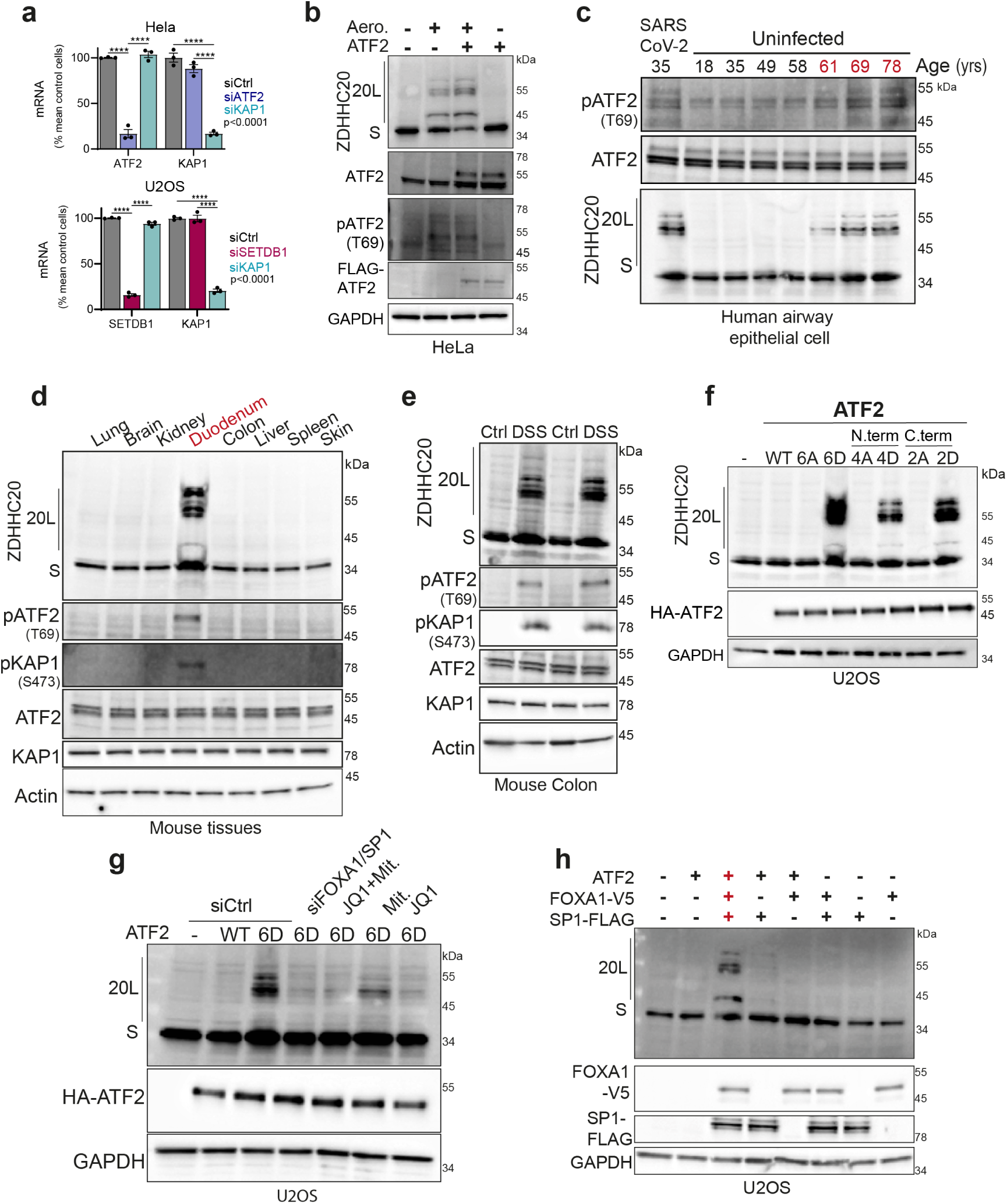
ATF2 and SETDB1-KAP1 dependent induction of ZDHHC20L expression. **a**, mRNA quantification validating efficiency of ATF2, KAP1 and SETDB1 siRNA depletion in U2OS and HeLa cells. **b–f,** Western blots (WB) for ZDHHC20 isoforms (Short-S, 20L), total/phospho-ATF2 in lysates from: b, HeLa cells overexpressing ATF2-FLAG (24 h), untreated or treated with proaerolysin toxin (10 ng/ml, 1 h at 37 °C, washed, further 8 h incubation); c, primary human airway epithelia (see Fig. 1b); d, mouse tissue extracts (as indicated); e, colon extracts from mice treated ± DSS (7 d) followed by 3 d recovery (d,e blots include total/phospho-KAP1); f, U2OS cells overexpressing ATF2 WT or indicated phospho-mutants (6A, 6D, 4A, 4D, 2A, 2D). **g,** WB as in f, in U2OS cells treated (24 h) with FOXA1 inhibitor (JQ1, 100 nM), SP1 inhibitor (mithramycin, 100 nM), or siRNA-depleted (72 h) for FOXA1 and SP1. **h,** WB analysis as in f, U2OS cells expressing ATF2-HA, SP1-FLAG, and FOXA1-V5, individually or combined.

### Activation of the transcription factor ATF2 triggers ZDHHC20L expression

To elucidate the molecular mechanisms underlying ZDHHC20L expression, we mined experimental gene occupancy data from ENCODE-3 (Encyclopedia of DNA Elements)^40,41^ hosted in the UCSC Genome Browser (Transcription Factor ChIP-seq Clusters)^42^. We previously found that ZDHHC20L is translated from upstream in-frame AUGs, generating longer transcripts. We also showed that FOXA1 and SP1 are transcription regulators, which bind approximately 5 kb upstream of the in-frame start codons, in HepG2, human liver cancer, cells^37^. Here, we found that the stress-regulated transcriptional activator ATF2^43^ was recruited ∼5 kB further upstream (Fig. 1c) and evaluated its role in ZDHHC20L expression. In Hela cells, where ZDHHC20L can be induced by aerolysin^37^, we found that siRNA-silencing of ATF2 (Supplementary Fig. 1a) significantly reduced the toxin-induced ZDHHC20L expression (Fig. 1d).

Different kinases activate ATF2 in response to stress or growth signals^44–50^ and aerolysin treatment also led to ATF2 (both endogenous and over-expressed) phosphorylation, on Thr-69, within its transactivation region^46^ (Fig. 1d, Supplementary Fig. 1b). The presence of multiple phospho-bands at around >55 kDa suggests phosphorylation on several residues and/or of various ATF2 isoforms^51^. ATF2 phosphorylation could be detected under all tested conditions leading to ZDHHC20L expression: in primary lung epithelial cells from older donors or subjected to SARS-CoV-2 infection (Supplementary Fig. 1c), in the duodenum of mice (where ZDHHC20L is constitutively expressed^37^ – Supplementary Fig. 1d), and in the colon of mice upon chemical-induced colitis (Supplementary Fig. 1e). Thus, ATF2 phosphorylation is consistently associated with ZDHHC20L expression.

To assess the importance of ATF2 phosphorylation in regulating ZDHHC20L expression, we generated ATF2 phospho-site mutants, either mimicking phosphorylation by aspartic acid substitution or preventing it by alanine substitution, targeting four residues in the N-terminal activation domain Thr-52, 69, 71 and 73 ^50^ (to D or A – coined 4D and 4A), two in the C-terminal regulatory domain Ser-490 and 498 ^44^ (2D and 2A), or all six combined (6D and 6A). Using bone-derived U2OS cells, which only express the canonical form of ZDHHC20, overexpression of any of the ATF2 phospho-mimetic mutants was sufficient to induce ZDHHC20L expression, in the absence of any stimulus (Fig. 1e, Supplementary Fig. 1f). This process still required FOXA1 and SP1, as it was blocked by the combined depletion or pharmacological inhibition of these transcription factors with JQ1 and mithramycin respectively (Supplementary Fig. 1g). Consistently, wild-type ATF2, but not ATF2-6A, when co-expressed with FOXA1 and SP1, also led to ZDHHC20L expression (Fig. 1e), which did not occur when only two of these factors were over-expressed (Supplementary Fig. 1h). To explore an additional functional outcome beyond controlling expression of ZDHHC20L during stress, we investigated whether ATF2 activation is protective against exposure to aerolysin, which like other pore-forming toxins disrupts plasma membrane integrity that can lead to cell death through various pathways^52,53^. Silencing ATF2 increased cell sensitivity to aerolysin (Fig. 1f), which was rescued by expressing wild type (WT) or the phosphomimetic 6D ATF2 but not the 6A mutant (Fig. 1f). Thus, activated ATF2 drives ZDHHC20L expression and has a protective effect towards aerolysin-induce cell death.

### *ZDHHC20L* is repressed by a heterochromatin-inducing complex

We further analyzed ENCODE ChIP-seq data and identified reported binding sites for the protein scaffold KAP1 (KRAB domain-associated protein 1) and the histone methyltransferase SETDB1 (SET Domain Bifurcated Histone Lysine Methyltransferase 1), near the ATF2 binding regions in U2OS and K562 lymphoblast cells (Fig. 1c) ^54,55^. These two proteins are part of a chromatin-modifying complex that mediates histone 3 lysine 9 trimethylation (H3K9m3), leading to the formation of heterochromatin^54^ and transcriptional repression^54–57^ (Fig. 1g, top panel). In U2OS cells, SETDB1 silencing (Supplementary Fig. 1a) alone induced ZDHHC20L expression (Fig. 1h), which was suppressed by wild-type SETDB1 but not by a catalytically inactive mutant (Fig. 1i), confirming the need for its enzymatic activity. KAP1 silencing (Supplementary Fig. 1a) or knockout in haploid HAP1 cells (Fig. 1j) similarly triggered ZDHHC20L expression (Fig. 1j) and the emergence of extended ZDHHC20 mRNA transcripts, as shown by qPCR (Fig. 1k)^37^. KAP1 and SETDB1 therefore repress ZDHHC20L expression, in contrast to ATF2 that is required for stress-induced ZDHHC20L expression.

Given the involvement of SETDB1 and KAP1, we analyzed ENCODE ChIP-seq data to assess whether histone modifications had been reported at the ZDHHC20 locus. In U2OS cells, repressive H3K9me3 marks were detected around the SETDB1, KAP1, and ATF2 peaks (Fig. 1c). Additional H3K9me3 marks within the first intron overlapped a second SETDB1 peak, potentially reinforcing ZDHHC20L silencing. Active promoter marks H3K27Ac and H3K4me3 (Fig. 1g, lower panel) were found flanking these repressive regions and co-localizing with both the short-canonical and long ZDHHC20 transcription start sites (Fig. 1c). The enhancer mark H3K4me1 (Fig. 1g) aligned with FOXA1 and SP1 ChIP-seq peaks in HepG2 cells (Fig. 1c), which constitutively express ZDHHC20L even without stimulation^37^.

Altogether, these data show that under homeostatic conditions the expression of ZDHHC20L isoforms is repressed by the heterochromatin-inducing KAP1/SETDB1 complex.

### ZNF354A recruits the repressive complex to the ZDHHC20L regulatory region

KAP1/SETDB1 complexes are generally tethered to DNA by KRAB-containing zinc finger proteins (KZFPs) (Fig. 1g), of which approximately 380 are encoded by the human genome^54,55,57^. To identify the KZFP responsible for docking KAP1 at the *ZDHHC20* locus, we mined KRABopedia, a published ChIP-seq dataset generated from HEK293T cells overexpressing HA-tagged forms of each KZFP individually^58,59^. The binding sites of several KZFPs map to the promoter region of ZDHHC20 (Fig. 2a, Supplementary Fig. 2a), a pattern that is not uncommon for this class of proteins^58,59^. In contrast, ZNF317, ZNF354A, ZNF677, and ZNF793 have binding sites within the “repression” region of the acyltransferase gene. ZNF354A binding sites co-localize with the SETDB1 and H3K9me3 peaks upstream and downstream of the *ZDHHC20* TSSs (Fig. 2a, Supplementary Fig. 2a). We silenced each of the four KZFPs individually in U2OS cells using mixes of siRNAs (Supplementary Fig. 2b). Only the ZNF354A knockdown led to the expression of ZDHHC20L (Fig. 2b), which could be reverted by the expression of ZNF354A-HA (Supplementary Fig. 2c). Importantly, silence of ZNF354A did not lead to a decrease of the mRNA of its paralogue ZNF354B, with which it shares 86% identity (Supplementary Fig. 2de), and consistently ZNF354B silencing did not trigger ZDHHC20L expression (Supplementary Fig. 2f).

**Figure 2.**
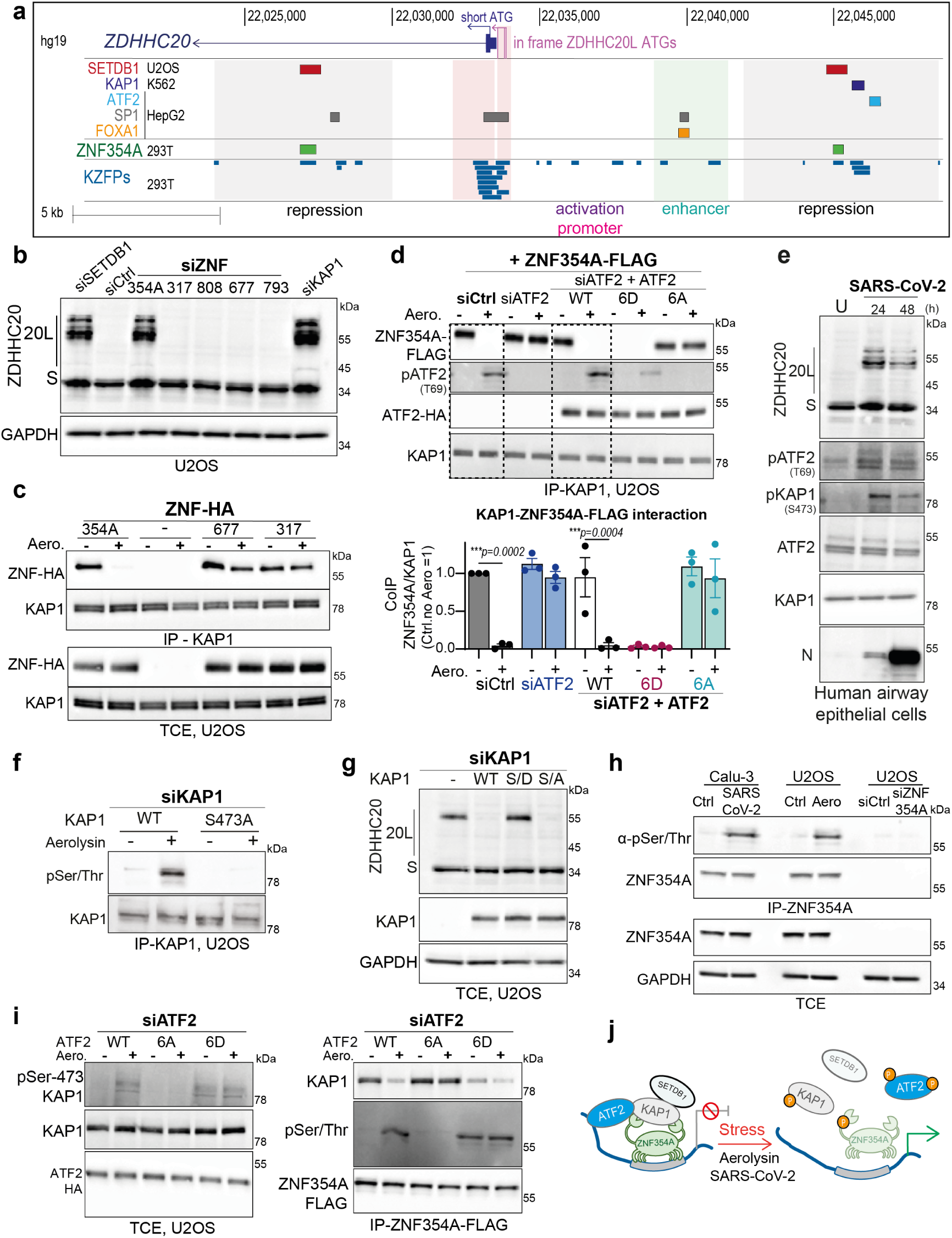
ZNF354A controls the expression of stress-induced ZDHHC20L. **a,** Overview of ZDHHC20 locus (intron, exon, 5′ UTR; hg19). Tracks: transcription start codons (ATGs), transcript annotations (GENCODE), ChIP-seq peaks (ENCODE: SETDB1/KAP1, ATF2, SP1, FOXA1), ZNF354A and KZFP peaks from HEK293T cells (squish view,) from KRABopedia^58^, https://tronoapps.epfl.ch/web/krabopedia/); **b,** Western blot (WB) of ZDHHC20 isoforms (Short-S, 20L) and GAPDH (loading control) in U2OS cells depleted of SETDB1, KAP1, or indicated KZFPs. **c,** Immunoprecipitation (IP) of endogenous KAP1 in U2OS cells overexpressing HA-tagged KZFPs, untreated or treated with proaerolysin toxin (10 ng/ml, 1 h at 37 °C, washed, incubated further 8 h). Blots: co-IP fractions and total cell extracts (TCE). **d,** IP-KAP1 as in (c) from siCtrl or siATF2 U2OS cells expressing ZNF354A-FLAG complemented with indicated ATF2 constructs (WT, 6D, 6A; 24 h). Results mean ± SEM; dots represent independent experiments (n = 3); P values vs. untreated, two-way ANOVA, Tukey’s correction. **e,** WB as (b) in human primary lung epithelial cells infected with SARS-CoV-2 (MOI 0.1). **f,** IP-KAP1 in siKAP1 U2OS cells expressing KAP1-WT or S473A mutant, treated with aerolysin as in (c). Blots show KAP1 and phospho-Ser/Thr proteins in IP fractions. **g,** WB of TCEs as (b) in siKAP1 U2OS cells expressing KAP1 WT, S473A (S/A), or S473D (S/D). **h,** IP-ZNF354A from Calu-3 cells infected with SARS-CoV-2 (24 h), aerolysin-treated U2OS cells (as in c), or siCtrl/siZNF354A-depleted U2OS cells. Blots show ZNF354A and phospho-Ser/Thr proteins (IP fractions) and ZNF354A and GAPDH (TCE). **i,** IP-ZNF354A –HA from U2OS cells expressing indicated ZNF354A-FLAG and ATF2 constructs, aerolysin-treated as (c). Blots: IP fractions (KAP1, ATF2, phospho-Ser/Thr); TCE (total/pS473-KAP1, ATF2-HA). **j,** Model of ATF2-KAP1-ZNF354A repressive complex before and after stress.

**Supplementary Figure 2.**
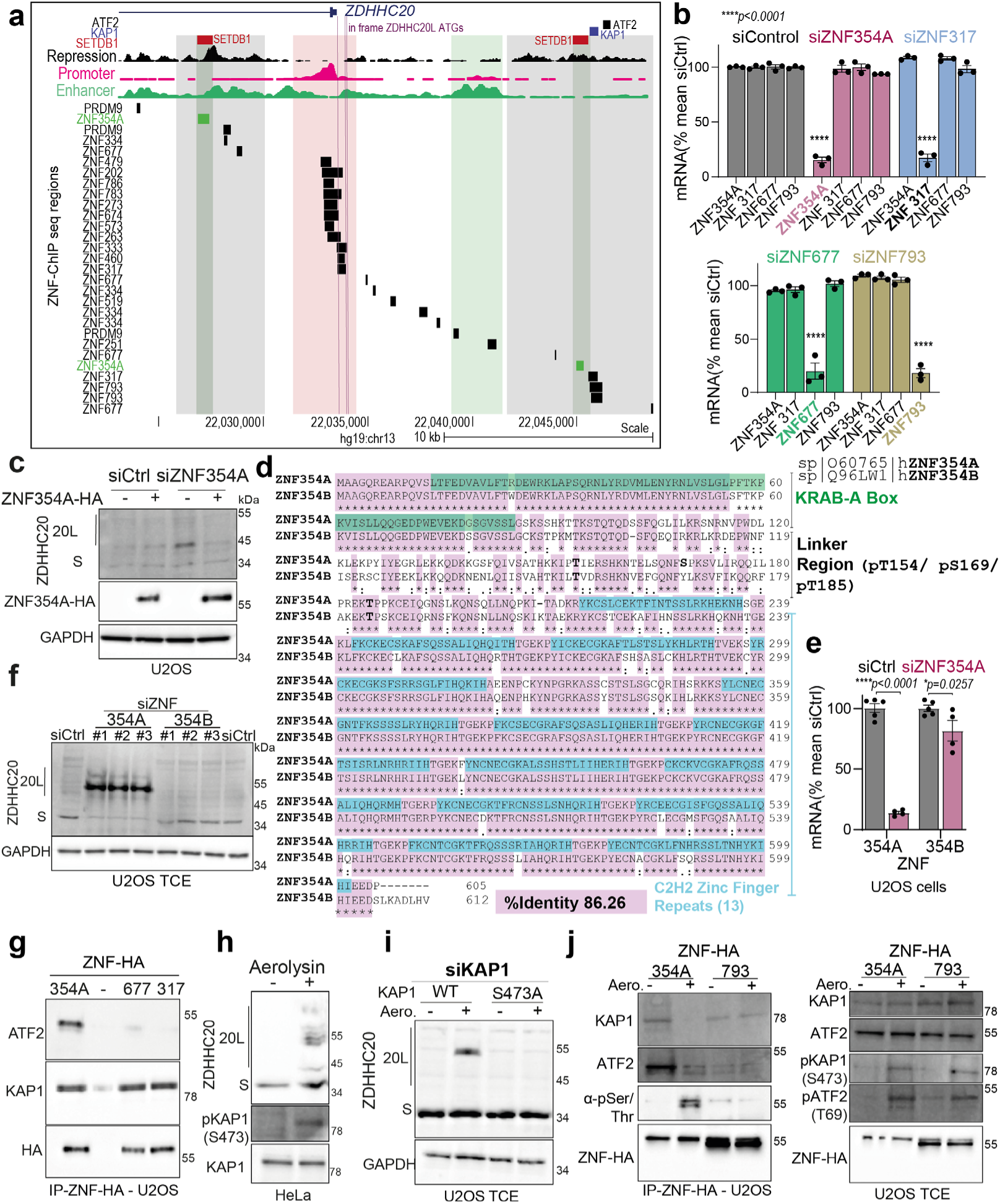
ZNF354A controls the expression of stress-induced ZDHHC20L. **a,** ZDHHC20 locus (hg19), indicating transcription start codons, transcripts (GENCODE), histone modifications, and ChIP-seq peaks SETDB1/KAP1/ATF2 (ENCODE); ZNF354A/KZFP, from HEK293T (KRABopedia^58^). See also fig 2a. **b,** mRNA levels of indicated KZFPs in U2OS cells after correspondent siRNA depletion (72 h). Mean ± SEM, each dot is an independent experiment n = 3; *p* values vs siCtrl, two-way ANOVA, Dunnett’s correction.**c**, Western blot (WB) of ZDHHC20 isoforms and GAPDH (loading control) in siZNF354A U2OS cells complemented with ZNF354A-HA. **d**, Protein sequence alignment of the indicated ZNF354A and ZNF354B Uniprot sequences with highlighted KRAB-A-Box (green), linker region with phosphorylated residues, 13 uniprot-annotated C2H2 Zinc-finger repeats and identity-residues (pink). **e**, mRNA levels as in b for ZNF354A and ZNF354B. Mean ± SEM, each dot is an independent biological replicate n = 5 siCtrl and n=4 siZNF354A; *p* values by two-way ANOVA, Tukey’s correction. **f**, WB as in c, in U2OS cells siZNF354A or siZNF354B depleted with three independent oligos. **g,** WB from IP-ZNF-HA fractions from U2OS cells expressing the indicated KZFP proteins. Blots show ATF2, KAP1 and ZNF-HA in IP-HA fractions. **h**, WB as in c, including total/phospho-KAP1 in HeLa cells ± proaerolysin toxin (10 ng/ml, 1 h at 37 °C, plus 8 h). **i**, WB as in c in siKAP1 U2OS cells expressing WT or S473A KAP1 mutant. **j**, IP-ZNF-HA from U2OS cells expressing indicated constructs, treated as in d. Blots show KAP1, ATF2, phospho-Ser/Thr (IP fractions); phospho-ATF2/KAP1 and total ZNF-HA (total cell lysates).

We next tested the specificity for ZNF354A versus other KZFPs by immunoprecipitation (IP) experiments. Upon IP of KAP1 from U2OS cells overexpressing ZNF354A or two irrelevant KZFPs, all three KZFPs co-precipitated with KAP1, as the presence of canonical KRAB domains would predict ^60^. Interestingly, the interaction between KAP1 and ZNF354A was disrupted by aerolysin treatment, which was not the case for the two other KZFPs (Fig. 2c). Thus, under basal conditions, ZNF354A recruits the KAP1 repressive complex to the 5′ regulatory region of DNA loci such as the ZDHHC20 locus. This interaction is disrupted upon aerolysin treatment, leading to the dissociation of ZNF354A and KAP1.

### Dynamics of the KAP1-ZNF354A containing repressive complex

Since ATF2, KAP1 and ZNF354A bind to the same region within the *ZDHHC20* 5’UTR, we tested whether they could be part of a single complex. We immunoprecipitated KAP1 from ZNF354A-FLAG-expressing cells, which were depleted or not of endogenous ATF2 using siRNA, and recomplemented with wild-type or ATF2 phosphorylation mutants (6A and 6D). Under resting conditions, KAP1 co-precipitated ZNF354A, as well as ATF2 (Fig. 2d). The co-precipitation between the three proteins was also observed when performing IPs against the ZNF354A HA tag (Supplementary Fig. 2g). Despite the disruption of the complex, ATF2 – especially its endogenous phosphorylated form–remained associated with KAP1 (Fig. 2d). These results indicate that a subset of ATF2-KAP1 interactions occurs independently of ZNF354A-mediated chromatin association. Since ATF2 has not previously been found to associate with KAP1-nucleated repressive complexes, we tested whether it also interacts with other KAP1–KZFP complexes. IP of ZNF317, ZNF677, and ZNF793 did not lead to the co-IP of ATF2 (Supplementary Fig. 2g, j). Thus, ATF2, KAP1 and ZNF354A are part of a somewhat non-canonical repressive complex. Interestingly, expression of the phospho-mimetic ATF2-6D mutant was sufficient to disrupt the KAP1–ZNF354A interaction even in the absence of aerolysin (Fig. 2d), indicating that ATF2 acts as an upstream, stress-sensitive regulator of the complex. The mechanisms underlying ATF2 recruitment to ZNF354A-specific KAP1 complexes remain to be elucidated.

Since ATF2 undergoes phosphorylation upon stress, we wondered whether the same occurs for KAP1 and ZNF354A. KAP1 is known to undergo phosphorylation on Ser-824 and Ser-473 in response to DNA damage^61–64,65^ and on Ser-473 under oxidative stress^66^. Using an anti-phospho-Ser-473-KAP1 antibody, we detected KAP1 Ser-473 phosphorylation during SARS-CoV-2 infection, concomitant with ATF2 activation (Fig. 2e). KAP1 Ser-473 phosphorylation was also observed upon aerolysin treatment (Supplementary Fig. 2h), detected in the duodenum of mice (Supplementary Fig. 1d), and in the colon of mice upon chemical-induced colitis (Supplementary Fig. 1e). To determine whether Ser-473 is the major phosphorylation site, we expressed WT KAP1 or the S473A phospho-deficient mutant in cells silenced for the endogenous protein and exposed them to aerolysin. Western blot analysis of KAP1 IP fractions, using anti-phospho-Ser/Thr antibodies, showed no signal for the S473A mutant (Fig. 2f), validating Ser-473 as the primary phosphorylation site under our conditions. Finally, we verified that Ser-473 phosphorylation was necessary to trigger ZDHHC20L expression upon aerolysin treatment (Supplementary Fig. 2i). As with ATF2, expression of a phospho-mimetic S473D KAP1 mutant was sufficient to trigger ZDHHC20L expression without any stimulus (Fig. 2g).

We next probed for the potential phosphorylation of ZNF354A. Aerolysin treatment and SARS-CoV-2 infection both induced serine/threonine phosphorylation of ZNF354A, detected by phospho-Ser/Thr antibodies in ZNF354A-IPs (Fig. 2h). This phosphorylated band corresponded to ZNF354A, as it was absent in ZNF354A-silenced cells (Fig. 2h). Aerolysin did not induce the phosphorylation of a control KZFP, ZNF793 (Supplementary Fig. 2j). Finally, aerolysin-induced phosphorylation of KAP1 and ZNF354A depended on ATF2 phosphorylation, which alone was sufficient to drive all these events (Fig. 2i).

Together, these findings demonstrate that ZDHHC20L expression is repressed by a complex comprising ATF2, SETDB1, KAP1, and ZNF354A. Upon stress, ATF2 is phosphorylated, followed by phosphorylation of KAP1 and ZNF354A, leading to dissociation of the KAP1– ZNF354A complex and induction of ZDHHC20L expression (Fig. 2j).

### p38 and JNK trigger phosphorylation of the repressive complex components

To further explore the regulatory mechanisms of the ATF2-KAP1-ZNF354A complex, we first identified the ZNF354A phosphorylation sites. Three residues, located in the linker region between the KAP1-binding KRAB domain and the DNA-binding domains, were found by phospho-proteomics in breast cancer (Ser-169 and Thr-185)^67^, and a subgroup of large T-cell lymphoma (T154)^68^ (https://www.phosphosite.org). This linker region is the most diverse between ZNF354A and ZNF354B. We generated the phospho-mutants of the three ZNF354A sites and expressed them in ZNF354A-silenced U2OS cells. A phospho-Ser/Thr signal was observed upon aerolysin treatment of WT ZNF354A-, but not S169A ZNF354A-expressing cells (Fig. 3a). The signal was reduced, but not absent, when expressing the T154A or T185A mutants (Fig. 3a), indicating a leading phosphorylation event at Ser169, as confirmed using phospho-tag gels (Fig. 3b). Ser-169 is absent in ZNF354B (Supplementary Fig. 2d).

**Figure 3.**
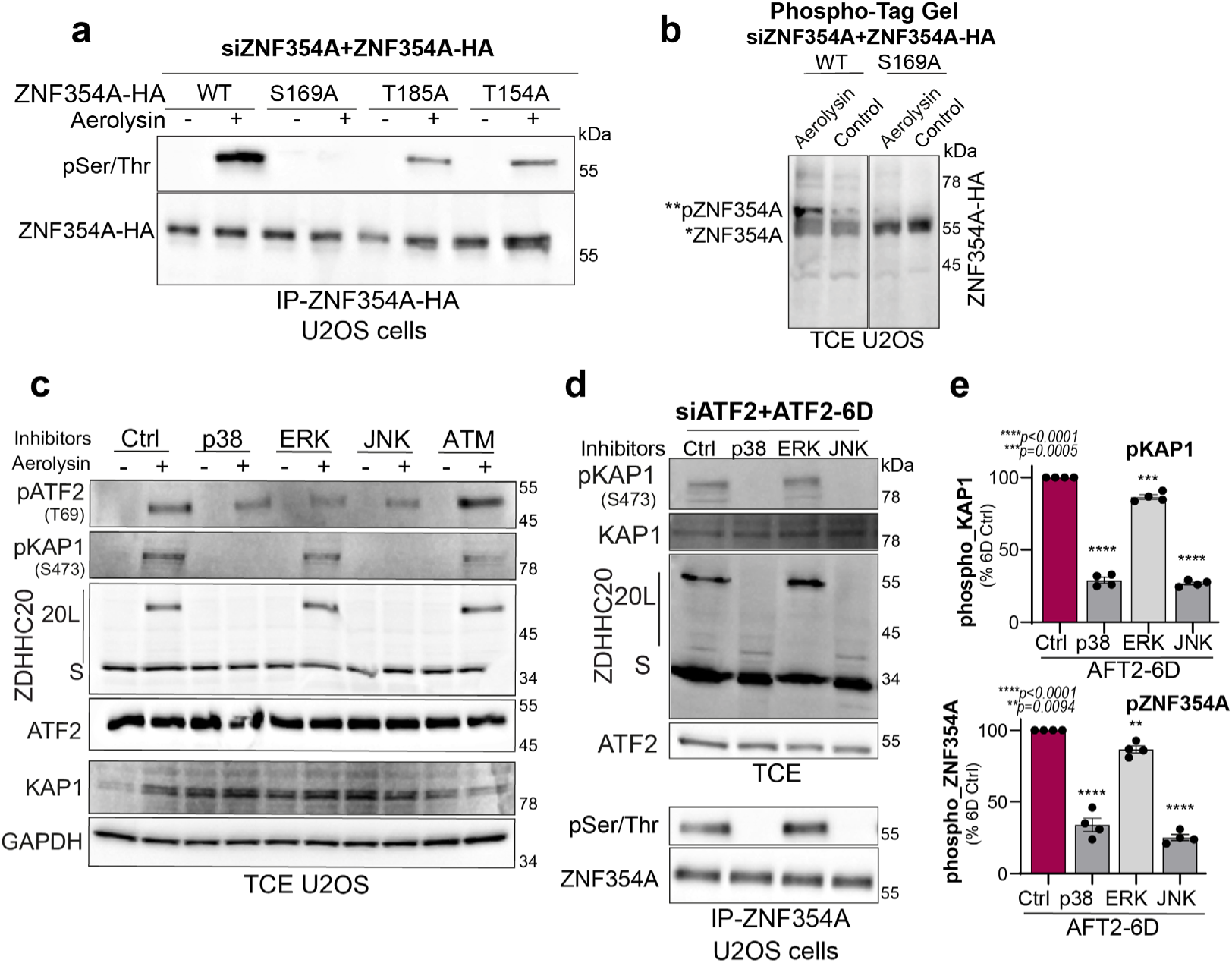
p38 and JNK phosphorylation of ZNF354A. **a,** Immunoprecipitation (IP) of ZNF354A-HA from U2OS cells (siZNF354A, 72 h) expressing WT or phospho-mutants of ZNF354A, untreated or treated with proaerolysin (10 ng/ml, 1 h at 37°C, washed, incubated additional 8 h). Blots show phospho-Ser/Thr proteins and ZNF354A in IP fractions. **b,** Phospho-Tag Western blot of total cell extracts (TCE) from U2OS cells treated as in (a). **c,** Western blot of ZDHHC20 (Short-S, 20L isoforms), phospho-KAP1 (S473), phospho-ATF2 (T69), total KAP1, ATF2, and GAPDH (loading control) in TCE from U2OS cells treated with proaerolysin as in (a), plus kinase inhibitors. **d,** Western blot as in (c) from TCE and IP-ZNF354A of U2OS cells (siATF2, 72 h) expressing ATF2-6D phosphomimetic mutant (24 h), treated with kinase inhibitors. **e,** Quantification of phospho-KAP1 and phospho-Ser/Thr-ZNF354A bands from blots in (d). Mean ± SEM; dots represent independent experiments (n=4). Statistical analysis by two-way ANOVA, Dunnett’s test; P values compare treated conditions to untreated control (Ctrl).

We next searched for the kinases involved in ZDHHC20L expression. We tested a panel of inhibitors targeting MAPKs (*e.g.* JNK, p38, and ERK) and ATM since these are known to modify KAP1 and ATF2 following different stresses, including infection^69,70^. Inhibition of either p38 or JNK abolished the aerolysin-induced expression of ZDHHC20L and KAP1 phosphorylation, and reduced phosphorylation of ATF2. ATF2 phosphorylation was also sensitive to ERK inhibition (Fig. 3c). p38 and JNK inhibitors similarly affected KAP1 and ZNF354A phosphorylation and ZDHHC20L expression in cells expressing the ATF2 6D mutant, in the absence of stressor (Fig. 3de). Thus, phosphorylation of ATF2, KAP1 (on Ser-473) and ZNF354 (on Ser-169) occurs in a p38– and JNK-dependent manner and is required for the expression of ZDHHC20L.

### ZNF354A determines the genetic specificity of the cellular response to oxidative stress

We next explored whether the de-repression by the ZNF354A-containing complex has a broader effect than just controlling ZDHHC20L expression and could be coupled with the activation of specific host stress response/repair pathways. To identify genes that reside in the vicinity of the 1,665 ZNF354A peaks, we analyzed our ChIP-seq data from HEK293T cells overexpressing HA-tagged ZNF354A^35,58^ using GREAT (Genomic Regions Enrichment Annotations) with a 300 kb window from the TSS as the cut-off for gene association ^71^. GREAT analysis mapped the ZNF354A peaks to the regulatory regions of 2,116 genes associated with a variety of ontologies (Fig. 4a). Among the enriched pathways, selenium metabolism stood out due to its connection to defense against ROS (Fig. 4a). Further manual curation of the gene list led to the identification of additional selenium-related genes (Supplementary Fig. 3a).

**Figure 4.**
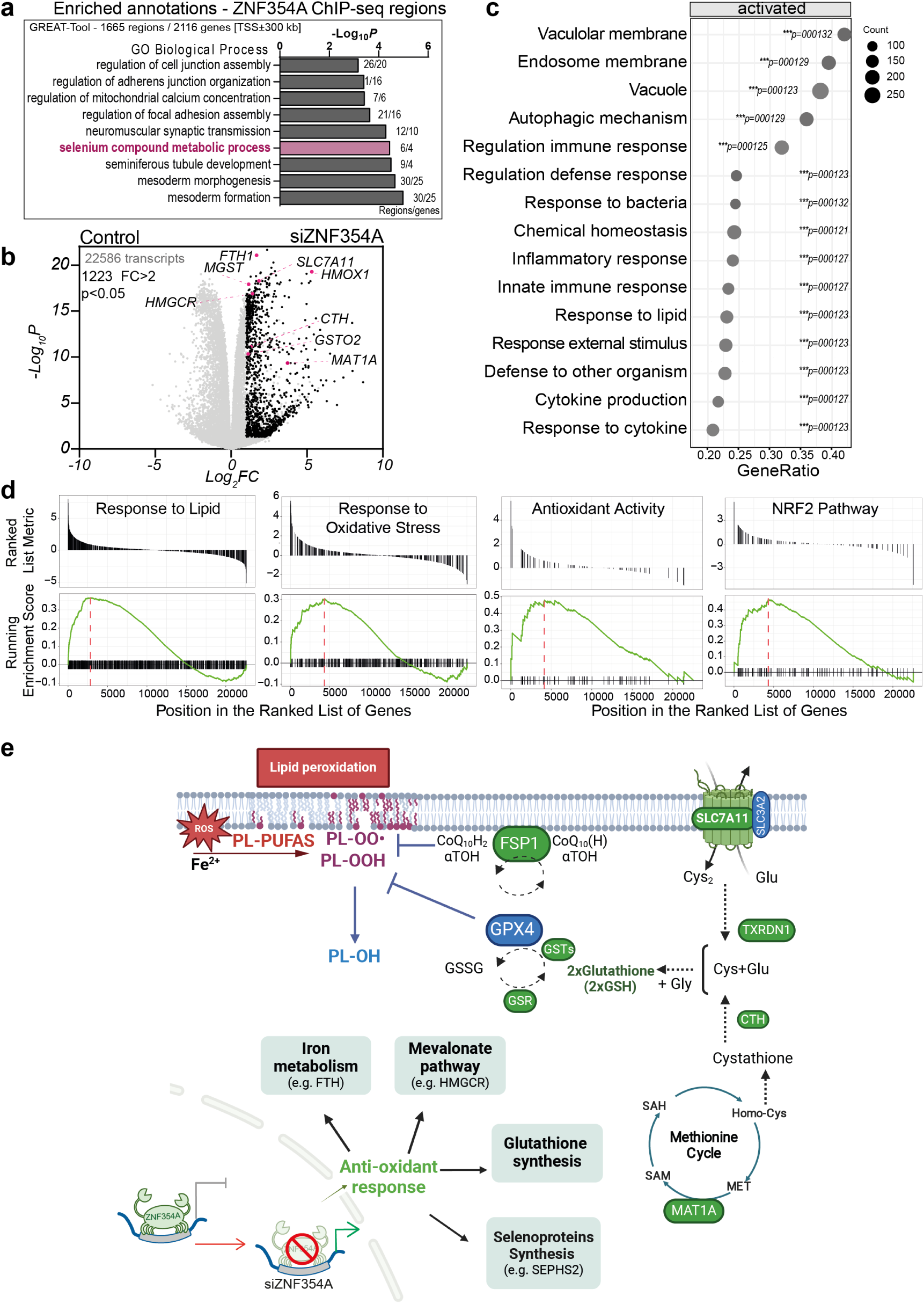
ZNF354A is a master regulator of antioxidant responses. **a,** Ontologies (GREAT analysis) from ZNF354A ChIP-seq peaks in HEK293T cells. Histogram shows enriched terms ranked by P value. **b,** Volcano plot of RNA-seq analysis from U2OS cells (siCtrl vs. siZNF354A, 72 h), displaying expression changes of 22586 transcripts. Log2 fold change (FC) vs. −log10 P value plotted. Highlighted genes involved in GSH synthesis (MAT1A, SLC7A11, GSTO2, MGST, CTH), iron metabolism (HMOX1, FTH), and mevalonate pathway (HMGCR). Black dots: significantly upregulated genes (FC > 2, P < 0.05). **c,** Gene Set Enrichment Analysis (GSEA) dot plot highlighting significantly enriched GO-terms in siZNF354A cells. Gene count, gene ratio, and specific values indicated. **d**, GSEA plots for Biological Process (BP) and Cellular Component (CC) GO categories. **e,** Model overview of ZNF354A-regulated selenium-/thiol-dependent and independent pathways in phospholipid peroxide detoxification. ZNF354A targets highlighted in green. Main proteins/pathways upregulated by ZNF354A depletion are indicated (see Supplementary Table 3). In thiol-dependent pathways GPX4 converts phospholipid hydroperoxides (PL-OOH) to alcohols (PL-OH), dependent on glutathione (GSH) synthesis via cystine uptake and reduction. The methionine cycle contributes with precursors. In parallel the NAD(P)H/FSP1/ubiquinone system reduces peroxyl radicals in phospholipids (PLOO•). SLC7A11 (xCT), cystine/glutamate antiporter; TXNRD1, thioredoxin reductase-1; CTH, γ-cystathionase; MAT1A, methionine adenosyltransferase; GSTs, GSH-S-transferases; GSR, GSH-disulfide reductase; GPX4, glutathione peroxidase-4; GSSG, oxidized glutathione; FSP1, ferroptosis suppressor protein; CoQ10H2, ubiquinol; CoQ10(H), ubiquinone; SEPHS2, selenophosphate synthetase-2; FTH, ferritin heavy chain-1; HMGCR, 3-hydroxy-3-methylglutaryl-CoA reductase. Created with BioRender.com. See methods or https://tronoapps.epfl.ch/web/krabopedia/ for full datasets

The identified redox-related genes included: *MAT1A* (methionine adenosyltransferase 1A), which synthesizes S-adenosylmethionine, a precursor for GSH biosynthesis^72^; *SLC7A11*, encoding the main subunit of the cystine/glutamate antiporter system x_c_^−^, essential for GSH production from extracellular cystine^73^; and *TXNRD1* (thioredoxin reductase-1 and its homolog *TXNRD3*), which can reduce cystine after its uptake by the x_c_^−^ system thereby contributing to GSH synthesis^74^. We also found 8 selenoprotein-encoding genes (Supplementary Fig. 3a). These proteins, of which the human proteome counts 25, contain a selenocysteine in their primary sequence which they use to reduce H₂O₂ and organic hydroperoxides^12,75^.

To test whether ZNF354A is indeed a regulator of antioxidant defense genes, we performed RNA-seq upon ZNF354A knockdown in U2OS (Supplementary Fig. 4a), which led to the identification of 1,223 significantly upregulated genes (fold change >2, p-adjusted 0.05) (Fig. 4b). Gene Set Enrichment Analysis (GSEA) indicated that these genes are involved in protective responses against external insults, including bacteria, viruses and chemical stresses, as well as antioxidant activity, lipid perturbations, and the NRF2 pathway (Fig. 4cd, Supplementary Fig. 4b, Supplementary Table 3, and Supplementary dataset), pointing in the same direction as the above ChIP-seq analysis of ZNF354A-overexpressing HEK293T cells (Supplementary Fig. 3a).

Because the NRF2 pathway has a well-established regulatory role in antioxidant responses and has been reported to be positively regulated by KAP1^76^, we tested its potential involvement in ZDHHC20L expression. While silencing ZNF354A leads to ZDHHC20L expression, this was prevented by the additional silencing of NRF2 (Supplementary Fig. 4cd). Thus, together with SP1 and FOXA1^37^, NRF2 also positively regulates ZDHHC20L expression.

To validate the gene set of Supplementary Fig. 3a, we performed quantitative PCR (qPCR). In ZNF354A-silenced cells, the mRNA levels of *MAT1A, SLC7A11, TXNRD1,* the redox regulator *GLRX* (Glutaredoxin-1), *SEPHS2* (selenophosphate synthetase) and *FSP1* were all increased (Supplementary Fig. 4e), consistent with elevated protein levels of MAT1A (Supplementary Fig. 4a). Complementary qPCR analysis in cells overexpressing ZNF354A for 24 h revealed downregulation of most of these target genes (Supplementary Fig. 4f), further supporting the regulatory role of ZNF354A.

As predicted by the analysis of the ZNF354A DNA binding regions (Fig. 4a), this KZFP is involved in the repression of genes encoding key antioxidant cellular defense effectors (Fig. 4, Supplementary Fig. 3a, 4b, and Supplementary Table 3). Many of its target genes are indeed involved in GSH biosynthesis, iron metabolism, the mevalonate pathway, and the NADPH-FSP1-CoQ10 oxidoreductase system, which protects cells from lipid radical-induced damage^21,22^ (Fig. 4e). These various independent analyses point towards ZNF354A as a key regulator of the genetic control of cellular antioxidant defenses, both selenium/thiol-dependent and –independent, and thereby the cellular sensitivity to ferroptosis^7,12,77^.

To assess the evolutionary conservation of the pathway, we extended our analysis to the mouse ortholog of ZNF354A, ZFP354A. Using a liftover strategy ^42,78^, we mapped human ZNF354A binding sites to the mouse genome assembly, identifying approximately 300 loci successfully converted. Remarkably, ZFP354A also binds near genes involved in antioxidant defense (Supplementary Fig. 3a). Consistently, silencing ZFP354A in mouse cells resulted in the expression of mouse ZDHHC20L (Supplementary Fig. 3b), mirroring our findings in human cells and supporting a conserved function of ZNF354A/ ZFP354A in regulating oxidative stress responses.

ZNF354A binds predominantly to transposable elements (TEs), with some additional binding observed in introns, as documented in Krabopedia ^58^ (Supplementary Fig. 3c). Among enriched TE families, ZNF354A displays a strong preference for LINE-1 (Long Interspersed Nuclear Elements)^15,19^. In the curated, redox-associated gene signature upregulated upon ZNF354A depletion, the nearest ZNF354A peaks were also largely associated with LINE-1 elements (Supplementary table 3). Future research will investigate how the emergence of ZNF354A and its cognate TE targets contributed to shaping the regulatory network governing the LORD pathway in placental mammals. This includes examining the role of ZNF354B, which, despite its strong similarity to ZNF354A, exhibits a distinct binding profile with significantly fewer peaks, suggesting functional divergence (Supplementary Fig. 3e).

**Supplementary Figure 3.**
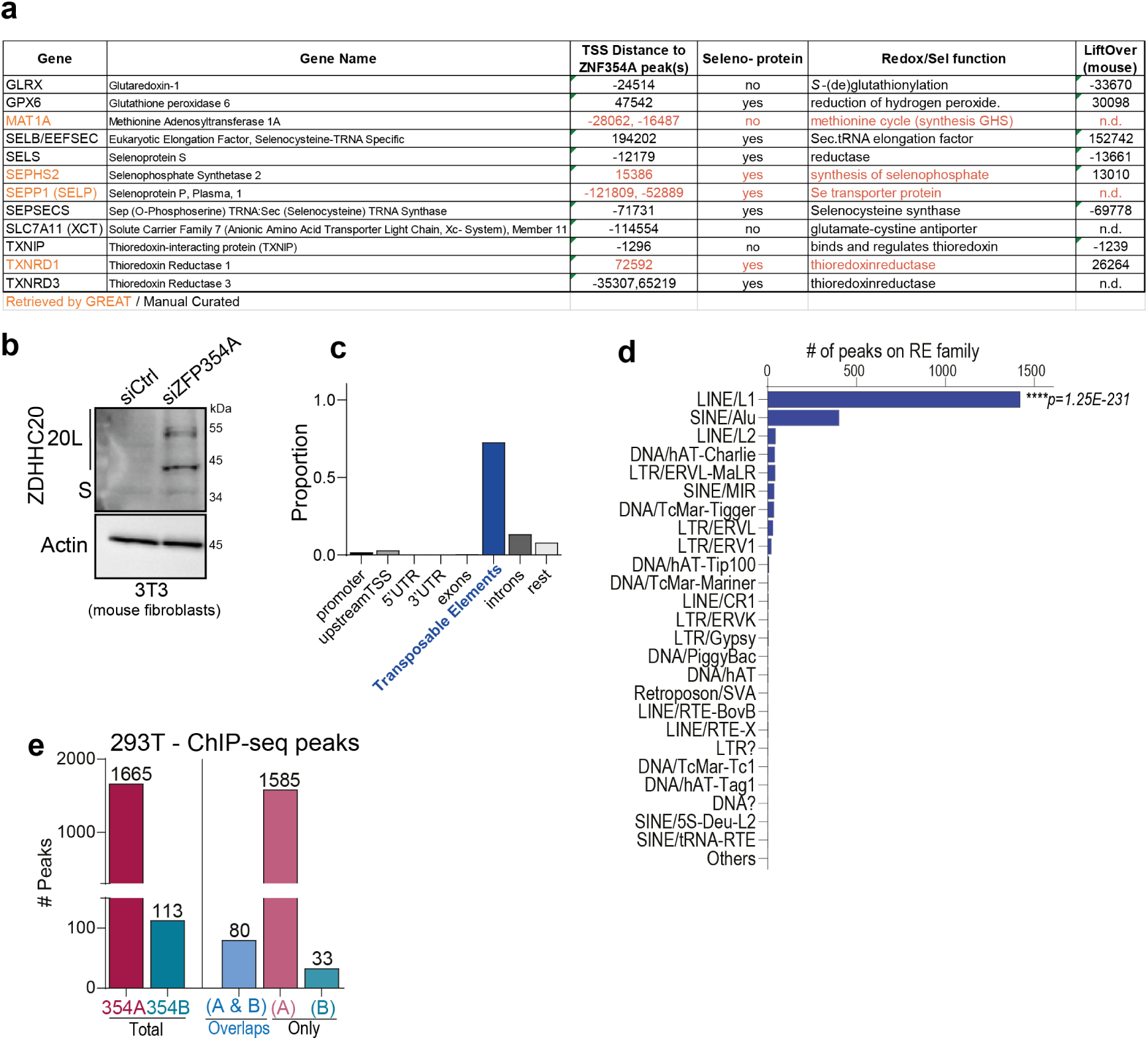
ZNF354A binds to preferentially to transposable elements and controls selenium-related genes. **a.** Selected genes associated with selenium pathways, antioxidant, and anti-ferroptotic activities identified near ZNF354A ChIP-seq peaks (within 300 kb from TSS) in HEK293T cells. Gene symbols, gene names, and distances to peaks are indicated. **b.** Western blots of 3T3 mouse fibroblasts transfected with siCtrl or siZNF354A (mouse ZFP354A). Blots show ZDHHC20 isoforms (Short-S 43 kDa and 20L 50 and 53 kDa) and loading controls (actin). **c.** Barplot depicting the distribution of peaks over genomic features with exclusive selection, meaning each peak is assigned to only one feature. **d.** Barplot showing the overlap of ChIP-seq peaks with TE families. P-values were computed using a binomial test. **e.** Comparison of ChIP-seq peaks for ZNF354A and ZNF354B, showing specific peaks and direct overlaps (≥1 bp). See methods or https://tronoapps.epfl.ch/web/krabopedia/ for full datasets.

**Supplementary Figure 4:**
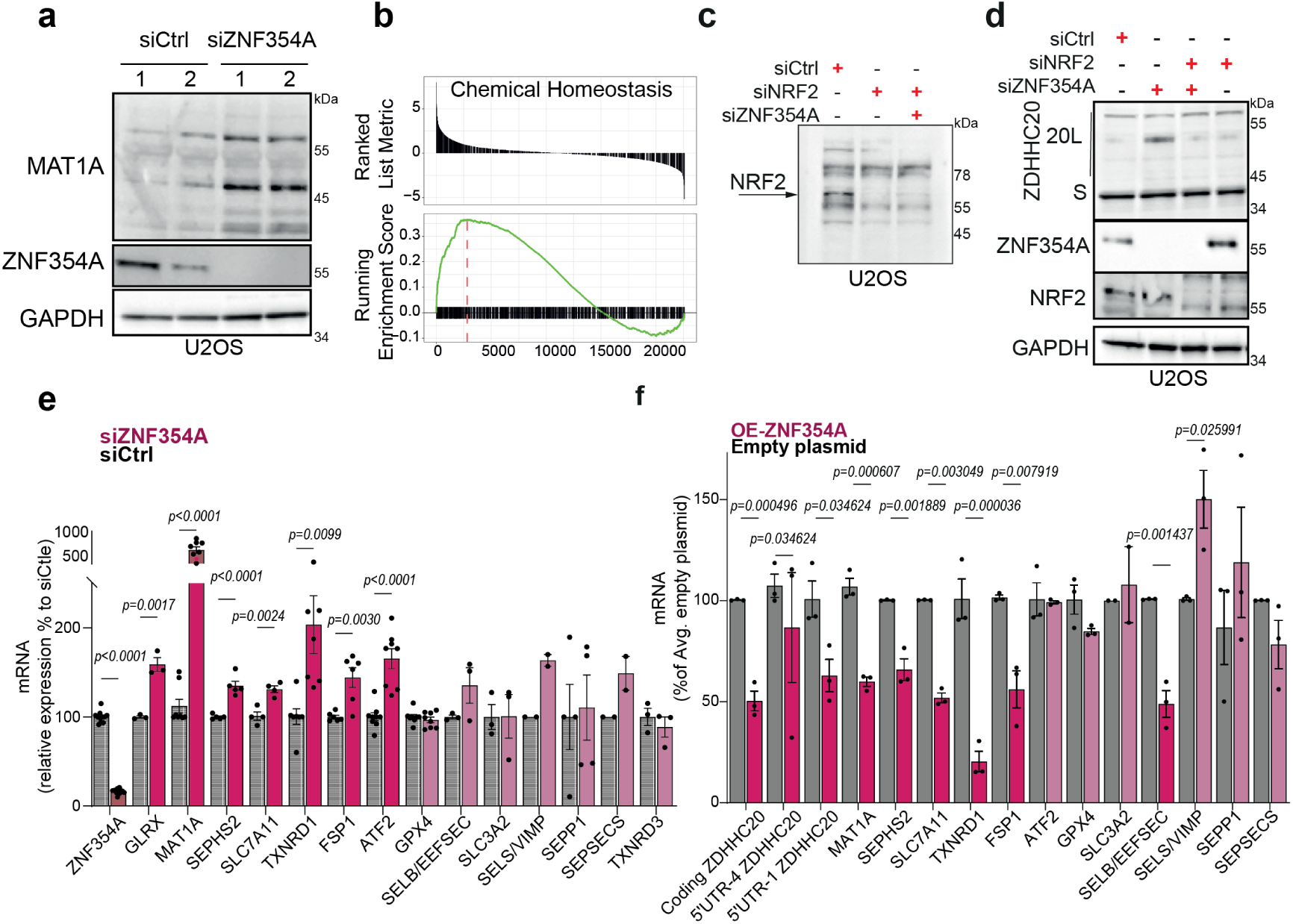
ZNF354A is a regulator of antioxidant responses. **a,** Western blot of MAT1A, ZNF354A and GAPDH (loading control) in TCE from U2OS cells treated 72h with 2 different RNAi control or against ZNF354A. **b,** GSEA plots of Chemical Homeostasis-Gene Ontology categories (Supplementary Table 2) from U2OS cells transfected with siCtrl or siZNF354A (72 h). **c,** mRNA quantification of indicated transcripts in U2OS cells transfected with siCtrl or siZNF354A (72 h). Mean ± SEM; each dot represents an independent experiment. Statistical analysis by two-way ANOVA, Sidak’s test; *p* values compare to siControl. **d,** mRNA quantification of indicated transcripts in U2OS cells transfected 24 hours with Ctrl or ZNF354A expressing plasmids. Mean ± SEM; each dot represents an independent experiment. Statistical analysis by two-way ANOVA, Sidak’s test; *p* values compare to empty plasmid. **e-f,** Western blots showing NRF2 (**f**), ZDHHC20 (Short-S, 20L isoforms), ZNF354A, and GAPDH (loading control) (**f**) in U2OS cells depleted of NRF2, ZNF354A, or both (siNRF2, siZNF354A, siNRF2/ZNF354A; 72 h).

### ZDHHC20L expression is induced by lipid peroxidation

Since ZNF354A regulates genes involved in cellular responses to oxidation, we explored whether oxidative damage could be the common primary trigger of the ZDHHC20L expression observed following SARS-CoV-2 infection, pore-forming toxin treatment, colitis, and aging (Fig. 1). When oxidative stress was induced by treating U2OS cells for 4-8 h with H₂O₂ at 50 µM, a significantly lower concentration than used to induce DNA damage ^14^, ZDHHC20L was expressed in a dose-dependent manner (Fig. 5a).

**Figure 5.**
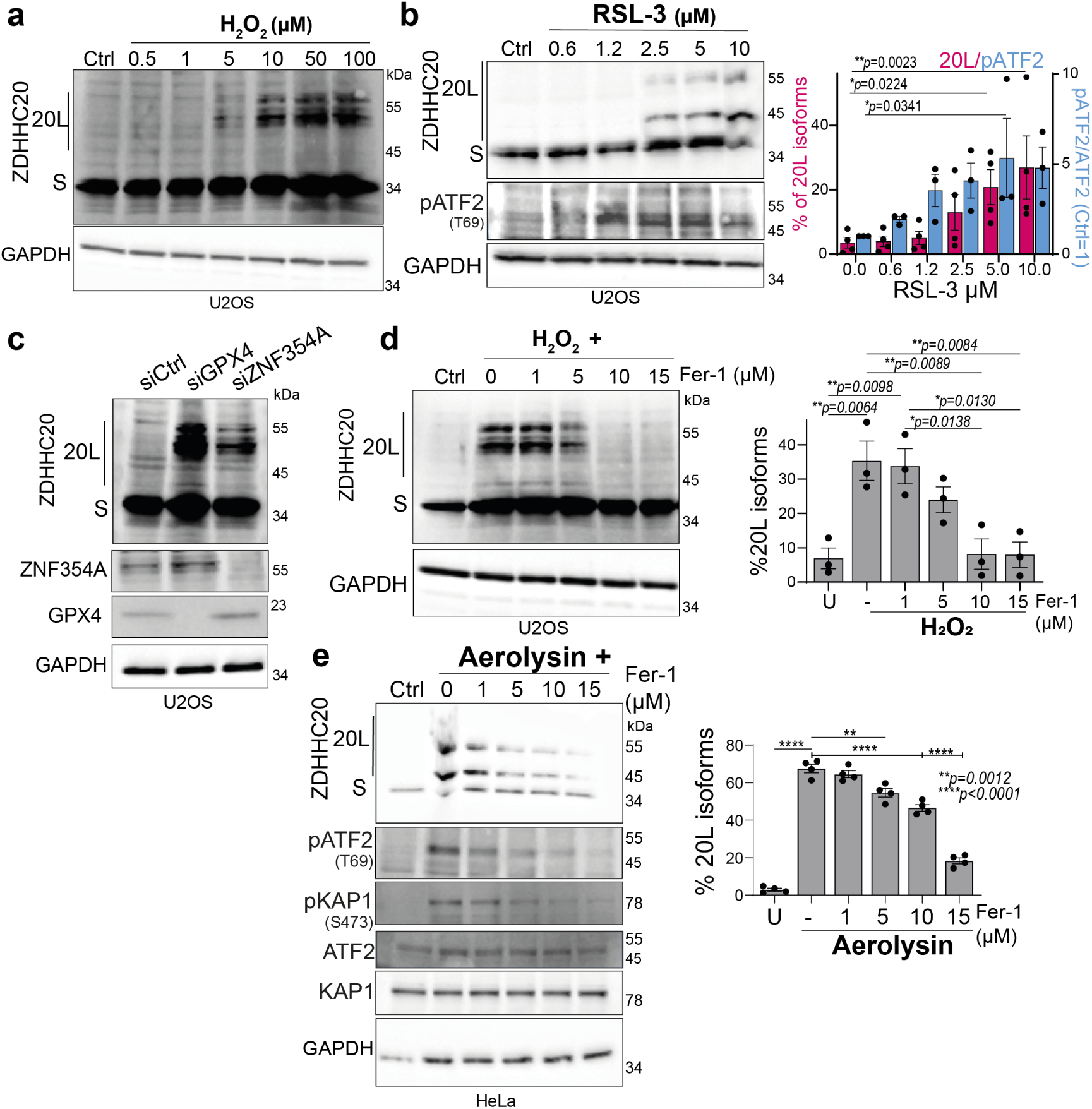
ZDHHC20L and ZNF354A respond to redox homeostasis and lipid peroxidation. **a–e,** Western blot (WB) analysis of ZDHHC20 (Short and 20L isoforms) and GAPDH (loading control) in U2OS cells treated as follows: increasing concentrations of H₂O₂ (8 h) (a), increasing concentrations of RSL3 (b), siControl, siGPX4 or siZNF354A (72 h) (c), H₂O₂ (50 μM) with increasing concentrations of ferrostatin-1 (Fer-1) (d), and HeLa cells treated with proaerolysin (10 ng/ml, 1 h, washed, incubated additional 8 h) (e). Graphs in (b,d,e) show percentages of 20L isoform and (b) ATF2 levels. Mean ± SEM; each dot represents an independent experiment. Statistical analysis by two-way ANOVA, Tukey’s test; *p* values as indicated.

**Supplementary Figure 5.**
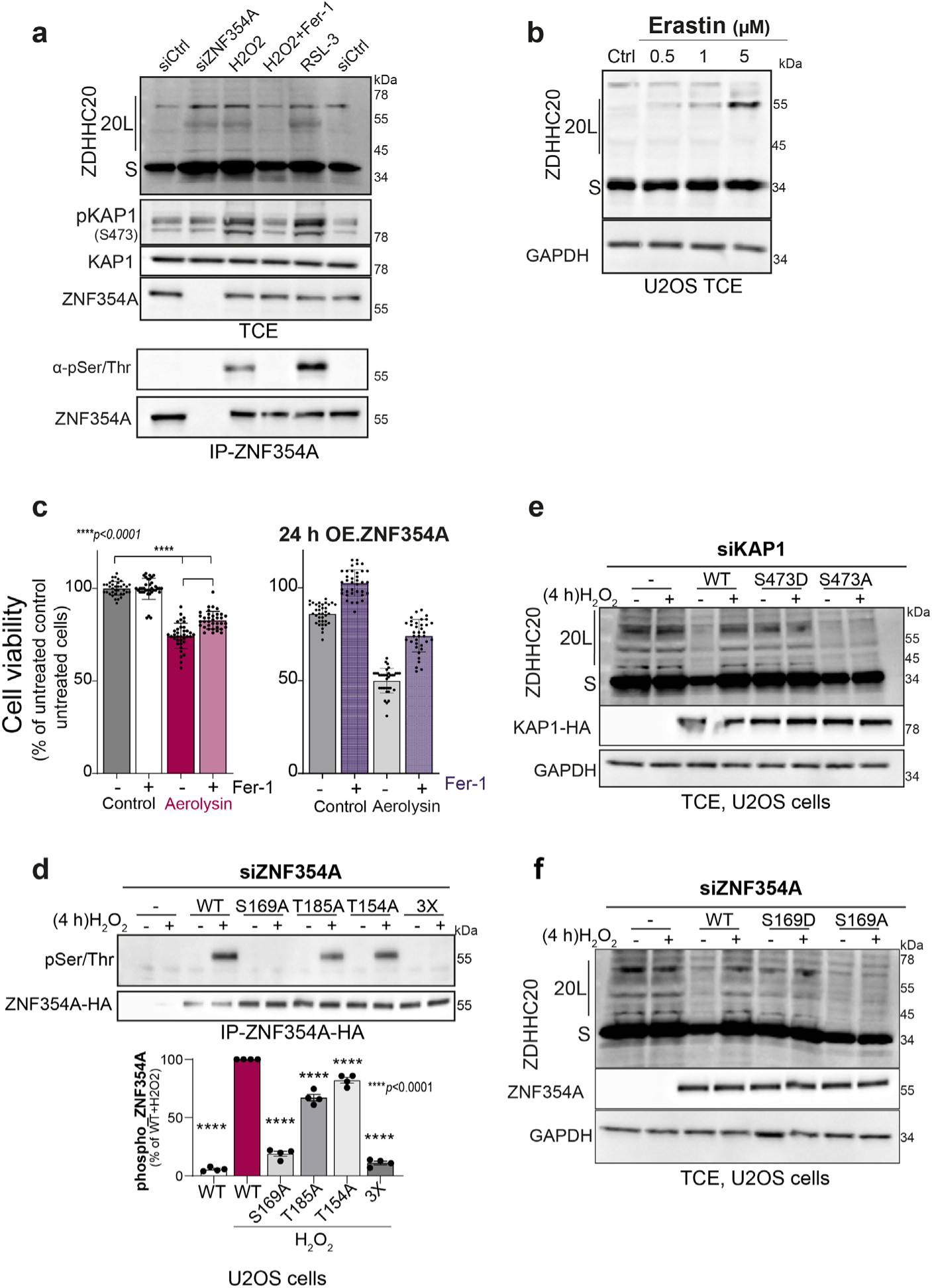
ZDHHC20L and ZNF354A respond to redox homeostasis and lipid peroxidation. **a,** Western blot (WB) of total cell extracts (TCE) and ZNF354A immunoprecipitates (IP) from U2OS cells (siCtrl or siZNF354A, 72 h) treated with H₂O₂ (50 μM, 7 h ± ferrostatin-1, Fer-1, 15 µM) or RSL3 (3 µM, 7 h). Blots show ZDHHC20 (20L and Short isoforms), phospho-KAP1, total KAP1, GAPDH (loading control), phospho-Ser/Thr proteins, and ZNF354A. **b,** WB (as in a) in TCE from U2OS cells treated with increasing concentrations of Erastin (16 h). **c,** Viability (ATP detection) of U2OS cells (Control or overexpressing ZNF354A, 24 h) ± Ferrostatin (15 μM) or/and proaerolysin (10 ng/ml, 1 h, washed, incubated additional 8 h last hours). Mean ± SD; representative assay (of three), n=36.; *p* values compare to control plasmid, by two-way ANOVA, Sidak’s test. **d**, IP-ZNF354A-HA in U2OS cells (siZNF354A, 72 h) overexpressing WT or phospho-mutant ZNF354A (3X mutant with all phospho-residues mutated to Ala), untreated or treated with H₂O₂ (50 μM, 4 h). Quantification of phospho-Ser/Thr/ZNF354A bands. Mean ± SEM; each dot represents an independent experiment (n=3). Statistical analysis by two-way ANOVA, Dunnett’s test; *p* values compare H₂O₂-treated WT. **e,** WB (as in a) in U2OS cells (siKAP1, 72 h) expressing WT KAP1 or indicated phospho-mutants, untreated or treated with H₂O₂ (as in d). **f**, WB (as in d) in siZNF354A, 72h cells expressing WT or indicated phospho-mutant ZNF354A, treated as in (d).

We next tested whether ZDHHC20L expression could be specifically triggered by inducing the accumulation of membrane lipid peroxides. These form when oxygen radicals interact with PUFA-containing phospholipids^5,13,79^. Under physiological conditions, their accumulation is avoided by the action of GPX4, the primary enzyme that detoxifies lipid peroxides, by reducing them to lipid alcohols^23^. Treating cells with the covalent GPX4 inhibitor RSL3 (RAS-selective-lethal-3)^25^ induced ZDHHC20L expression in a dose-dependent manner (Fig. 5b). H₂O₂ and RSL3 treatment also triggered phosphorylation of ATF2 (Fig. 5b), KAP-1, and ZNF354A (Supplementary Fig. 5a). Silencing of GPX4 similarly led to ZDHHC20L expression (Fig. 5c). Finally, since GPX4 requires GSH for its reduction, a third way of impairing its activity is to block cystine uptake via SLC7A11 (system Xc^-^). Erastin, a system Xc⁻ inhibitor ^80^, also induced ZDHHC20L expression in a dose-dependent manner (Supplementary Fig. 5b).

Upon induction of lipid peroxidation, detoxification can also be performed chemically using lipophilic peroxyl radical-trapping molecules such as Ferrostatin-1 (Fer-1)^81^. Ferrostatin-1 inhibited the H₂O₂-induced expression of ZDHHC20L in a dose-dependent manner (Fig. 5d). Importantly, ferrostatin also inhibited aerolysin-induced ATF2 and KAP1 phosphorylation, ZDHHC20L expression (Fig. 5e), and diminished aerolysin-induced cell death (Supplementary Fig. 5c). Altogether, these observations demonstrate that lipid peroxide accumulation induces ZDHHC20L expression, whereas inhibition of lipid peroxidation by Ferrostatin-1 prevents this induction.

We next verified whether H₂O₂ could recapitulate the events triggered by aerolysin exposure or SARS-CoV-2 infection. H₂O₂ treatment led to the phosphorylation of ATF2, KAP1 (on Ser-473), and ZNF354A (on Ser-169), in a ferrostatin-dependent manner (Supplementary Fig. 5a). H₂O₂-induced ZDHHC20L expression was prevented by the expression of the phospho-deficient KAP1 S473A and ZNF354A S169A mutants (Supplementary Fig. 5d-f). The H₂O₂-induced ATF2, KAP1 and ZNF354A phosphorylation and ZDHHC20L expression were p38– and JNK-dependent (Fig. 6ab). Inhibition of ERK reduced ATF2 phosphorylation moderately, diminished ZNF354A phosphorylation, but did not affect KAP1 phosphorylation or ZDHHC20L expression (Fig. 6ab). ATM inhibition mildly reduced phosphorylation of all three components and the expression of ZDHHC20L (Fig. 6ab). Finally, H_2_O_2_ treatment triggered the detachment of KAP1 from ZNF354A, while phosphorylated ATF2 interacted with KAP1 (Fig. 6c).

**Figure 6.**
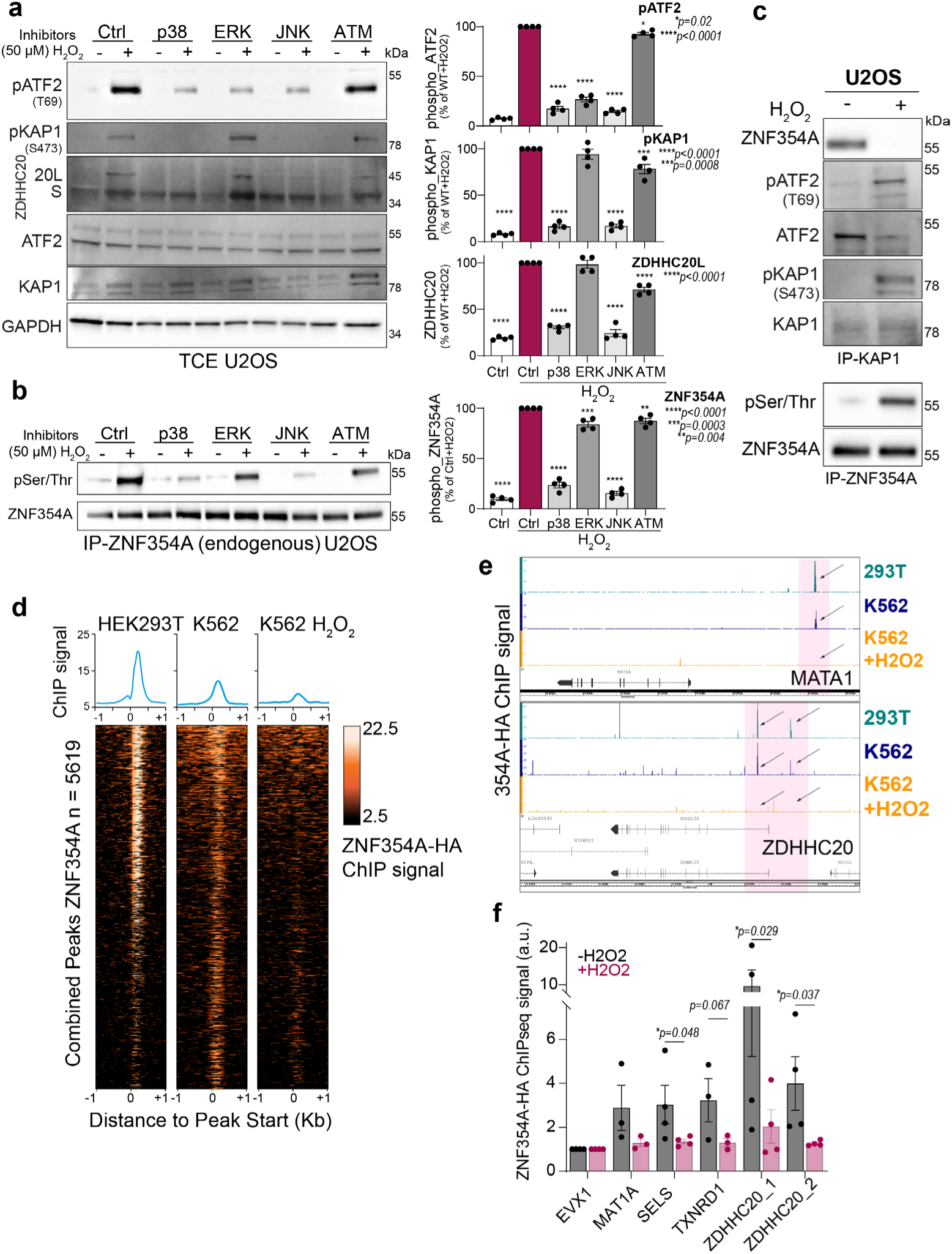
H₂O₂-dependent regulation of the ATF2-KAP1-ZNF354A epigenetic complex. **a,b**, Western blots of U2OS cells treated with H₂O₂ (50 μM, 4 h) in presence of indicated kinase inhibitors. Blots show ZDHHC20 (Short-S, 20L isoforms), phospho-KAP1 (S473), phospho-ATF2 (T69), total KAP1 and ATF2, GAPDH (loading control) (a), and phospho-Ser/Thr proteins with total ZNF354A (b). Quantification of phospho-ATF2, phospho-KAP1, ZDHHC20L (a), and phospho-Ser/Thr/ZNF354A (b) bands. Mean ± SEM; each dot represents an independent experiment (n=4). Statistical analysis by two-way ANOVA, Dunnett’s test; *p* values compare treated conditions to untreated control (Ctrl). **c,** Western blots as in (a), in KAP1 or ZNF354A immunoprecipitates from H₂O₂-treated cells as in (a). **d,** Heat maps of ChIP-seq profiles centered on ZNF354A-HA peaks (±1 kb) showing enrichment in HEK293T and K562 cells under untreated and H₂O₂-treated conditions (50 μM, 4 h). **e,** IGV browser screenshots illustrating selected ZNF354A ChIP-seq peaks (arrows) within *MAT1A*, and *ZDHHC20* loci from cells treated as described in (d). **f.** ChIP–qPCR quantification of ZNF354A-HA peaks at indicated gene regions (two peaks for *ZDHHC20*; *EVX1* used as control). Results mean ± SEM; dots represent independent experiments (n = 3). *p* values vs untreated (−H₂O₂), two-way ANOVA, Tukey’s correction.

**Supplementary Figure 6.**
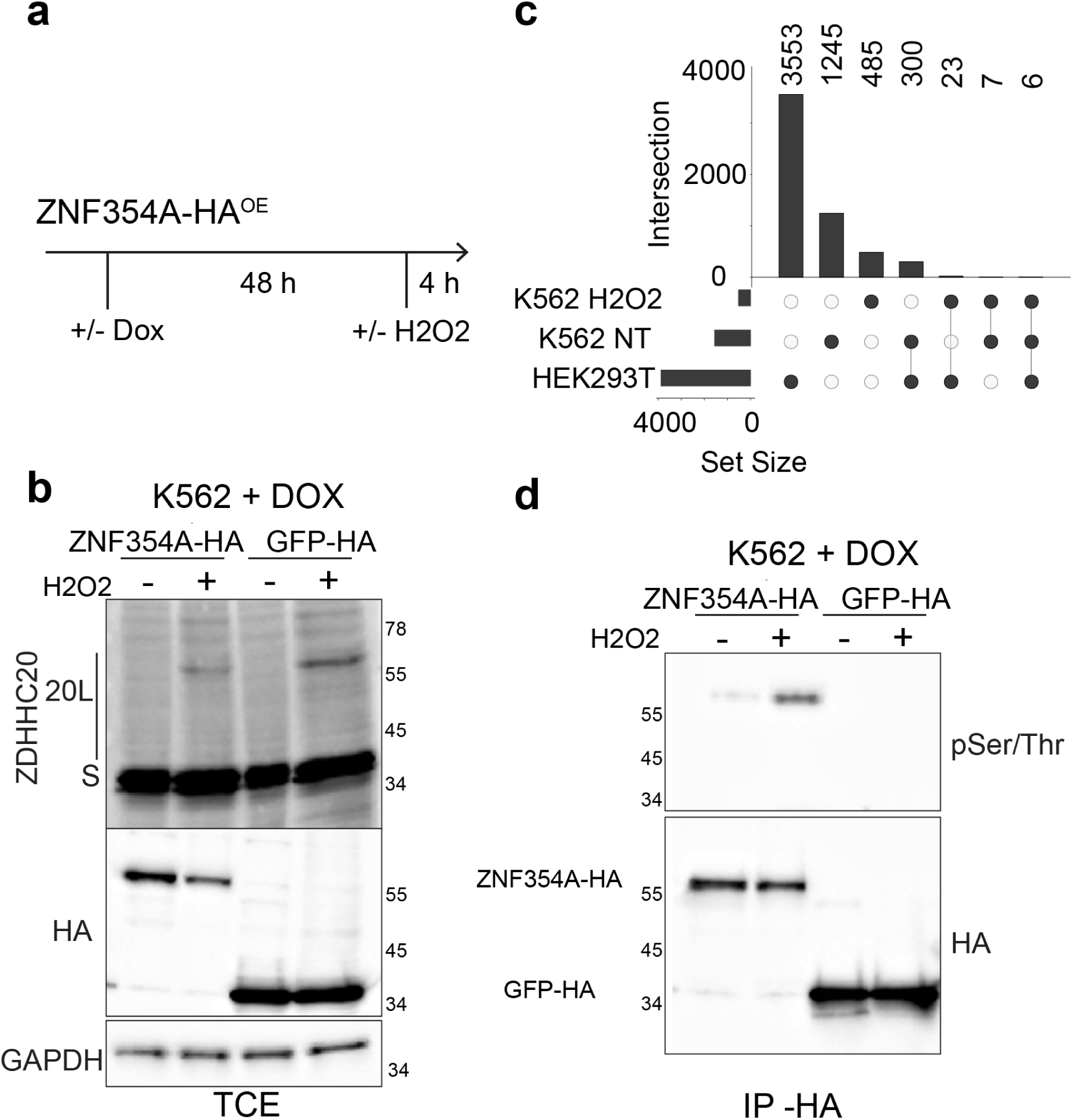
H₂O₂-dependent regulation of the ATF2-KAP1-ZNF354A epigenetic complex. **a,** Schematic of ChIP-seq experimental setup using K562 cells stably expressing control GFP-HA or ZNF354A-HA. ZNF354A expression was induced with doxycycline (1 μg/ml, 48 h) and verified by western blot (WB). Cells were untreated or treated with H₂O₂ (50 μM, 4 h). **b**, WB analysis of total cell extracts (TCE) showing ZDHHC20 (Short and 20L isoforms), HA-tagged protein expression, and GAPDH (loading control). HA immunoprecipitates (IP-HA) from K562 cells (a) were analyzed for HA-tagged proteins and phospho-Ser/Thr proteins. **c,** UpSet plot showing overlaps of ZNF354A ChIP-seq peaks identified in HEK293T cells (HEK293T), untreated K562 cells (K562_NT), and H₂O₂-treated K562 cells (K562_H₂O₂; 50 μM, 4 h). Horizontal bars indicate total peaks per condition; vertical bars represent intersections of shared peaks between conditions.

Altogether, these observations demonstrate that H₂O₂ recapitulates the observations made with aerolysin, and can trigger the expression of ZDHHC20L through the p38/JNK-dependent phosphorylation of ATF2, KAP1 (Ser-473) and ZNF354A (Ser-169), and the disassembly of the repressive complex.

### Stress-induced release of ZNF354A from regulatory loci

Thus far, our data indicate that the repressive complex contains ATF2-SETDB1-KAP1-ZNF354A and that upon lipid peroxidation-induced phosphorylation, KAP1 detaches from ZNF354A. We next tested whether, under these conditions, ZNF354A is released from the DNA. We performed genome-wide profiling of ZNF354A binding upon its overexpression in K562 cells, treated or not with H₂O₂ for 4 hours (Supplementary Fig. 6ab). As a preamble, we examined the genomic recruitment of this ZNF354A in untreated cells, and found it to be consistent with that previously recorded in 293T cells, with 20% of K562 peaks (306/1558) already identified in this other cell line (Supplementary Fig. 6c), representing a significant overlap compared to randomized regions (permutation test: 100 permutations, empirical *P*-value = 0) (Fig. 6d). Additionally, the ZNF354A-HA signal intensity over peaks was highly consistent between the ChIP-seq experiments in the two cell lines (Fig. 6d). H_2_O_2_ treatment induced a severe drop in the number of ZNF354A-HA peaks (521 vs. 1558 peaks), and a significantly lower overlap with the peaks previously identified in HEK293T (6% vs. 20%, Fisher’s exact test, *P*-value = 2.465e-16) (Fig. 6d). Moreover, ZNF354A occupancy was significantly reduced at the majority of identified peaks (Fig. 6e, Supplementary Fig. 6c). That ZNF354A detaches from the DNA sites upon H₂O₂ treatment was further confirmed by ChIP-RT-qPCR for specific peaks associated with selected redox target genes, including *MAT1A*, *SELS*, *TXNRD1* and *ZDHHC20* (Fig. 6f). Together, these findings demonstrate that H₂O₂ treatment leads to the chromatin eviction of ZNF354A, to allow expression of previously repressed genes, notably those involved in antioxidant response.

### ZNF354A acts as a regulator of ferroptosis sensitivity

To test the functional relevance of ZNF354A in controlling an antioxidant gene program, we tested whether silencing ZNF354A expression would protect cells from lipid peroxidation-induced death. We first monitored lipid peroxidation, using two different fluorescent probes – C11-BODIPY and LiperFluo, the latter specifically detecting oxidized phospholipids^82,83^. Silencing ZNF354A expression led to a decrease in lipid peroxide levels (Fig. 7ab). ZNF354A-depleted cells also showed a moderate reduction in lipid peroxide accumulation following GPX4 inhibition with RSL3 compared to controls (Fig. 7ab). This suggests that in the absence of ZNF354A, cells have an increased ability to prevent the formation or detoxify lipid peroxides. Consistently, staining with the membrane-impermeant dye propidium iodide (PI) indicated that plasma membrane integrity was more preserved in ZNF354A-depleted cells than in control cells (Fig. 7c). Finally, cell viability was monitored by measuring ATP levels, following RSL3 or H₂O₂ treatment. Depletion of ZNF354A by siRNA led to a marked increase in viability in response to both RSL3 and H₂O₂ (Fig. 7d), as well as the cystine import inhibitors Erastin (Supplementary Fig. 7a).

**Figure 7.**
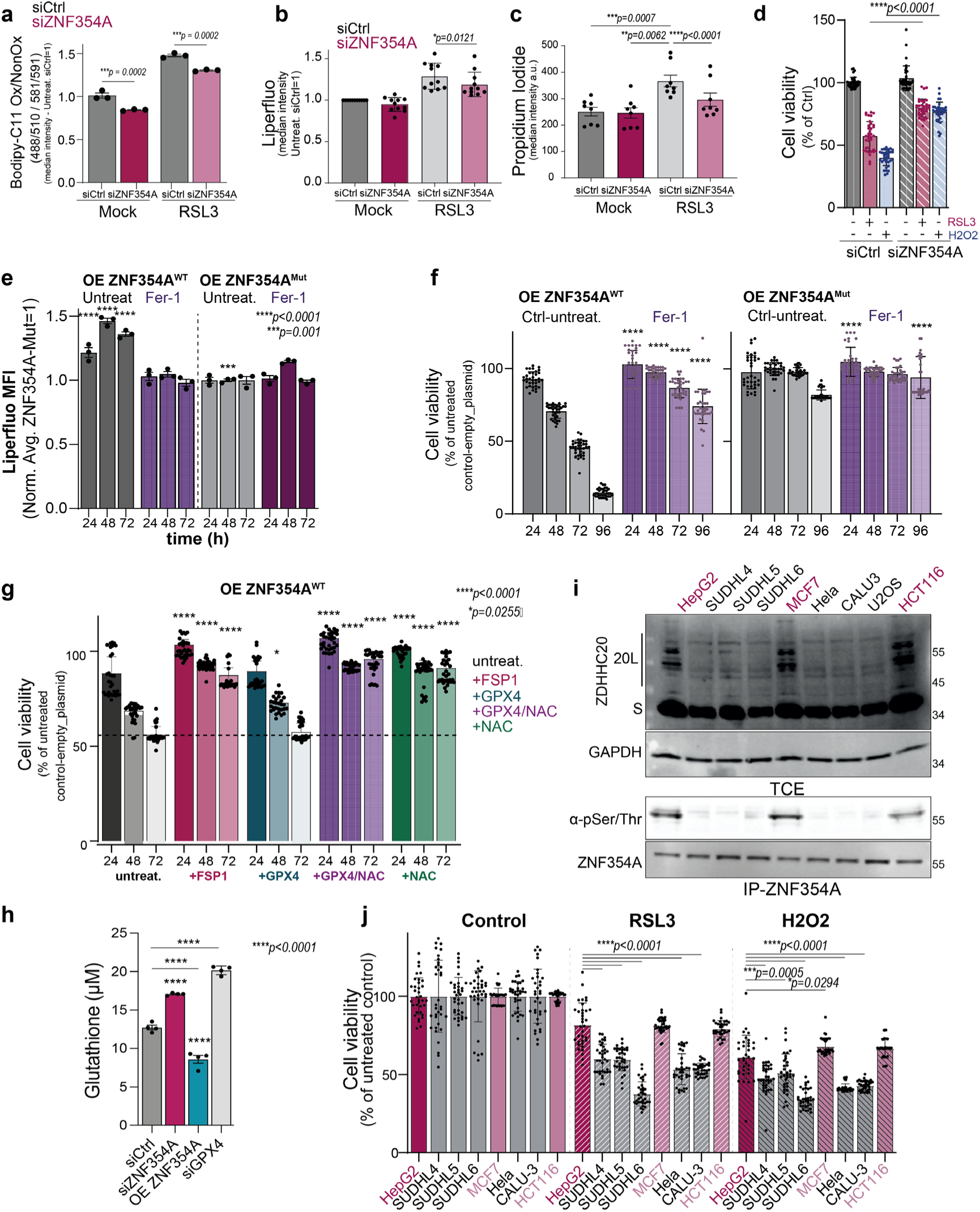
ZNF354A controls lipid peroxide detoxification and sensitivity to ferroptosis. **a–c**, Lipid peroxides measured by flow cytometry (Bodipy-C11, Liperfluo) and membrane permeability (propidium iodide) in U2OS cells transfected with siControl or siZNF354A (72 h) ± RSL3 (3 μM, 24 h). Mean ± SEM; dots indicate independent experiments.; *p* values compare to siControl, by two-way ANOVA, Tukey’s test. **d**, Viability (ATP detection) of U2OS cells (siControl or siZNF354A, 72 h) ± RSL3 (3 μM) or H₂O₂ (50 µM, 16 h). Mean ± SD; representative assay (of three), n=36.; *p* values compare to siControl, by two-way ANOVA, Sidak’s test. **e,f,** Lipid peroxidation (**e**) and viability (**f**) of U2OS cells expressing wild-type (WT) or KRAB-binding mutant ZNF354A ± ferrostatin-1 (Fer-1, 15 µM, added at 0 and 48 h post-transfection). Mean ± SEM and dots represent independent experiments (e). Mean ± SD normalized to untreated controls (empty plasmid) of representative assay (of three) n=33 (f). *p* values compare untreated controls vs Fer-1, by two-way ANOVA, Sidak’s test. **g**, Viability (as in f) of HeLa cells expressing ZNF354A, supplemented with FSP1, GPX4, GPX4 + N-Acetyl-Cysteine (NAC, 1 mM), or NAC alone. Mean ± SD; representative assay (of three), n=33.; *p* values compare untreated controls at each time point, by two-way ANOVA, Sidak’s test. **h**, The cellular level of GSH was measured in control cells and upon silencing of ZNF354A or GPX4 and upon over expression of ZNF354A. Mean ± SEM. **i**, Western blot of total cell extracts (TCE) and ZNF354A immunoprecipitates (IP) from indicated cancer cells, showing Short-S (42 kDa), 20L isoforms (∼49, 62, 69 kDa), GAPDH (loading control), total ZNF354A, and phospho-Ser/Thr proteins. **j**, Viability (as in d) of cancer cells (from h) ± RSL3 (3 μM) or H₂O₂ (50 µM, 16 h). Viability normalized to untreated controls (100%). Mean ± SD; representative assay (n=35). *p* values compare each condition to HepG2 reference, by two-way ANOVA, Dunnett’s test.

**Supplementary Figure 7.**
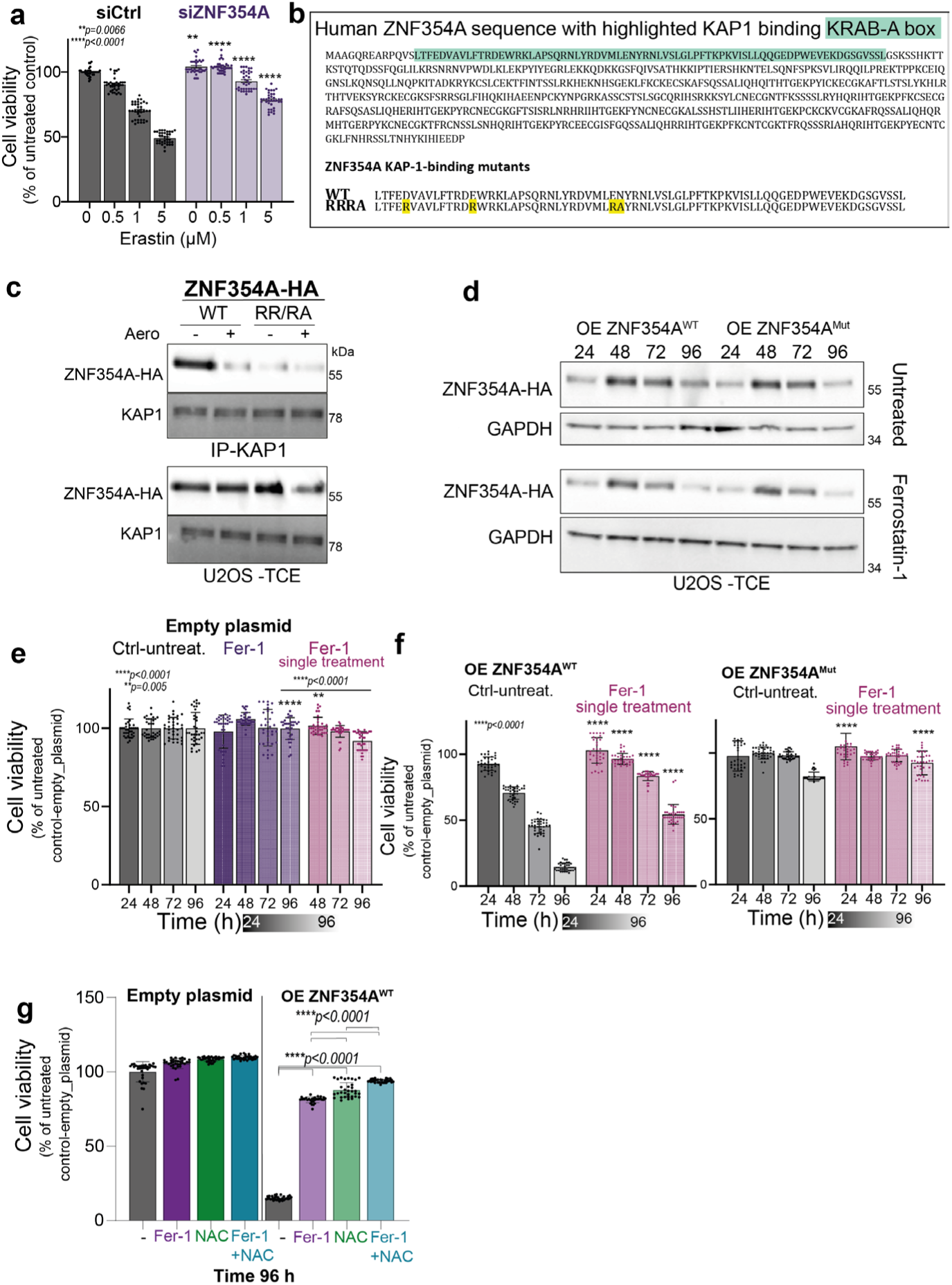
ZNF354A controls lipid peroxide detoxification and sensitivity to ferroptosis. **a**, Viability (ATP detection) of U2OS cells transfected with siControl or siZNF354A (72 h), treated with increasing concentrations of Erastin (16 h). Mean ± SD; representative assay (of three), n=36. Statistical analysis by two-way ANOVA, Sidak’s test; P values compare siControl vs siZNF354A. **b**, ZNF354A protein sequence with KAP1-binding KRAB-A box (green) and KAP1-binding mutantations. **c**, Immunoprecipitation (IP) of endogenous KAP1 from U2OS cells overexpressing wild-type (WT) HA-tagged ZNF354A or KAP1-binding mutant (RR/RA), untreated or treated with proaerolysin (10 ng/ml, 1 h, 37 °C), followed by washing and additional incubation (8 h, 37 °C). Western blots show co-immunoprecipitated proteins (CoIP) and total cell extracts (TCE). **d–f**, U2OS cells overexpressing WT or KRAB-binding mutant ZNF354A, treated ± ferrostatin-1 (Fer-1, 15 µM) at time 0 and 48 h post-transfection. d, Western blot analysis of TCEs showing ZNF354A-HA and GAPDH. **e,f**, Cell viability (ATP detection) in U2OS cells transfected with empty plasmid (e) or ZNF354A WT/KRAB-binding mutant (**f**), treated ± Fer-1 (single treatment, 15 µM at time 0). Results normalized to untreated empty plasmid controls (set as 100%). Mean ± SD; representative assay (of three), n=35. Statistical analysis by two-way ANOVA, Sidak’s test; p values compare untreated controls vs Fer-1 at each time point. **g**, Cell viability (ATP detection) in U2OS cells transfected with empty plasmid (e) or ZNF354A WT, treated ± Fer-1 (everyday treatment, 15 µM), or NAC (1mM everyday) or both treatments together. Results normalized to untreated empty plasmid controls (set as 100%). Mean ± SD; representative assay (of three), n=36. Statistical analysis by two-way ANOVA, Sidak’s test; p values compare untreated vs treated.

These observations show that silencing ZNF354A expression reduces the cellular sensitivity to ferroptosis. We next tested whether ZNF354A overexpression would sensitize them. We designed inactive ZNF354A mutants by mutating the ZFP-KRAB-A box of ZNF354A, a well-conserved interaction domain of KZFP with KAP1 (Supplementary Fig. 7b) ^60^, composed in particular of D18, E27, N45 and E46 ^60,84,85^. We generated a quadruple ZNF354A mutant construct with arginine substitutions of D18 and E27, or arginine and alanine substitutions of N45 and E46 (Supplementary Fig. 7b). While the mutant was expressed at WT levels, as predicted it failed to interact with KAP1 (Supplementary Fig. 7c). Overexpression of ZNF354A^WT^, but not ZNF354A^Mut^, led to an increase in oxidized phospholipids (Fig. 7e, Supplementary Fig. 7d), associated with significant cell death starting at 48 hours (Fig. 7f). Cell viability could be rescued by ferrostatin-1, to levels that depended on the frequency of Fer-1 treatment (Fig. 7ef, Supplementary Fig. 7e-g). When ZNF354A was overexpressed to levels that led to 85% cell death after 4 days, daily administration of Fer-1 reduced death to 22±16% (Supplementary Fig. 7g). Comparable protection was observed by treating cells with N-acetylcysteine amide (NAC), a cell-permeable cysteine derivative that restores GSH production, which reduced cell death to 12±5% (Supplementary Fig. 7g). The treatment with both Fer-1 and NAC reduced cell death to 7±1% (Supplementary Fig. 7g). Thus, cell death induced by ZNF354A overexpression is predominantly triggered by lipophilic ROS (≈75% contribution), with a minor contribution (<20%) by cytosolic or other ROS. Altogether, these observations show that ZNF354A silencing prevents excessive lipid peroxide accumulation and protects from cell death, while overexpression does the opposite.

As mentioned earlier, lipid peroxide detoxification relies mostly on GPX4 and FSP1, the latter operating in a GSH-independent manner via the NAD(P)H/FSP1/CoQ10 axis. We tested whether cell death induced by ZNF354A overexpression could be rescued by co-overexpression with either GPX4 or FSP1. FSP1 restored the viability of ZNF354A-overexpressing cells, consistent with the effect of Fer-1, while GPX4 did not (Fig. 7g). As this might reflect insufficient levels of GSH, we supplemented cells with NAC, which was sufficient to rescue ZNF354A-induced cell death, indicating that cellular GPX4 levels were not limiting (Fig. 7g). We then measured GSH levels to test whether they are influenced by ZNF354A. Silencing ZNF354A increased cellular GSH levels, as did to a greater extent silencing of GPX4 expression (Fig. 7h). Conversely, overexpression of ZNF354A led to a decrease in GSH (Fig. 7h). Thus, ZNF354A appears to act as a rheostat of cellular GSH levels. The induction of ferroptosis is of major therapeutic interest to kill cancer cells^25^. To probe the relevance of the ZNF354A pathway in predicting and controlling the sensitivity of tumor cell lines to ferroptosis, we first monitored the phosphorylation of ZNF354A. Only the cells expressing ZDHHC20L, i.e., HEPG2, MCF7, and HCT116 (Fig. 1a, 7i), showed phosphorylated ZNF354A, despite similar ZNF354A expression levels in all cells (Fig. 7i). We next submitted all cell lines to H₂O₂ or RSL3 treatment and monitored cell viability. Consistent with our hypothesis, HEPG2, MCF7, and HCT116 cells were more resistant to RSL3 and H₂O₂, compared to all other cancer lines (Fig. 7j).

Altogether, these findings indicate that ZNF354A regulates cellular GSH levels and lipid detoxification capacity, thereby modulating cellular sensitivity to lipid peroxidation-induced cell death pathways, in particular ferroptosis. In the context of cancer, the presence of phospho-ZNF354A or ZDHHC20L therefore appears predictive of tumor cell sensitivity to ferroptosis-inducing drugs.

### ZDHHC20L is an effector of the ZNF354A-mediated pathway

In this study, we have so far used the expression of ZDHHC20L as a proxy to decipher the ZNF354A-mediated pathway. We initially identified ZDHHC20L in SARS-CoV-2-infected cells and could show that ZDHHC20L greatly outperforms the canonical enzyme in modifying the envelope glycoprotein Spike^37,38^. We postulated that the virus had hijacked a stress response pathway—presumably activated to fight infection—for its own benefit. Here, we hypothesized that ZDHHC20L may act as a bona fide effector of the ZNF354A-mediated pathway, with a role in redox homeostasis. To explore this possibility, we sought to identify potential redox-regulatory substrates of ZDHHC20L. Given the central role of GPX4 in detoxifying lipid peroxides and protecting against ferroptosis, we hypothesized that ZDHHC20L might regulate GPX4 through this modification. During the course of our study, two reports were published indicating that S-acylation of GPX4 is critical for its function in controlling cellular sensitivity to ferroptosis ^86,87^. The first study showed that GPX4 undergoes acylation by ZDHHC20 on Cys-66^86^, while the second proposed acylation by ZDHHC8 on Cys-75^87^. Our screen of the 23 human ZDHHC enzymes, first tested in mixes of siRNA^38^, showed that only Mix 3 led to a marked decrease in GPX4 ^3^H-palmitate incorporation (Fig. 8ab). The enzymes targeted by Mix 3, namely ZDHHC9, 14, 18, and 20, were subsequently tested individually. Only ZDHHC20 modified GPX4 in our cellular system (Fig. 8cd). We further confirmed S-acylation using a click chemistry-based approach, so-called Click Acyl-PEG assay ^88^, which involves labeling cells with a clickable palmitate analogue, and subsequently labeling the added acyl chains *in vitro* with azide-PEG to induce a band shift detected by SDS-PAGE and Western blotting. A clear GPX4 band shift towards two upper bands could be observed in control HEK293T cells, which was absent in the corresponding ZDHHC20 KO line (Fig. 8e). The double band shift is indicative of acylation on more than one cysteine, suggesting that both reported cysteines at positions 66 and 75 could be simultaneously subject to S-acylation^86,87^.

**Figure 8:**
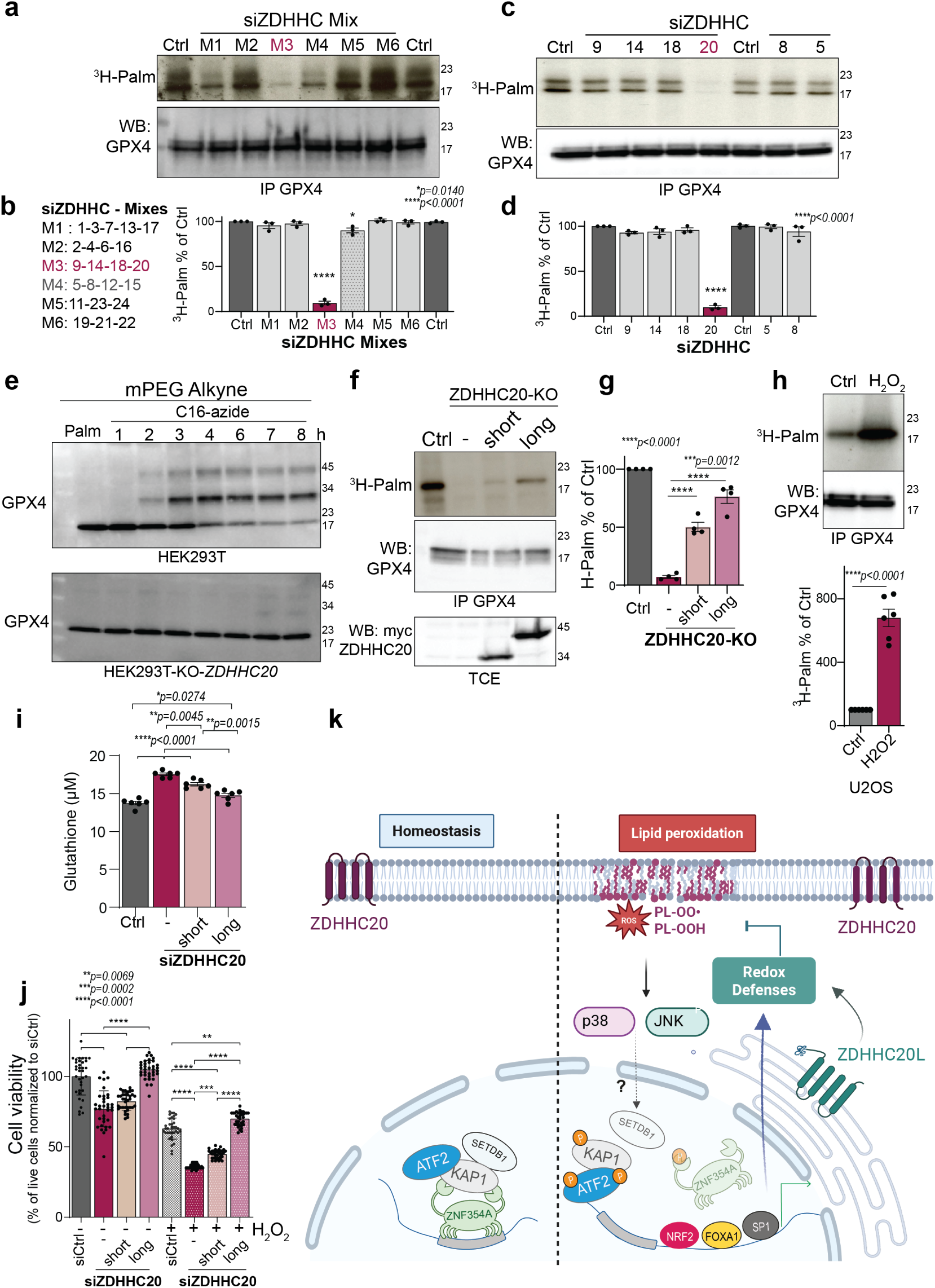
ZDHHC20L is an effector of the LORD pathway. **a–d.** Incorporation of ^3^H-palmitic acid (^3^H-palm) in GPX4 from U2OS cells silenced for 72 h with pooled (a) or individual (c) ZDHHC siRNAs. Cells were metabolically labeled for 3 h at 37 °C with ^3^H-palm; GPX4 was immunoprecipitated (IP-GPX4), resolved by SDS–PAGE, and analyzed by Western blot (WB) and autoradiography (^3^H-palm). The levels of ^3^H-palm incorporation in control RNAi (Ctrl) were set to 100% and results are mean ± SEM, and each dot represents an independent experiment (n=3). *p* values compare siCtrl, two-way ANOVA, Dunnett’s test. **e.** HEK WT or ZDHHC20-KO cells were incubated with 100 μM C16:0-azide for the indicated times or with palmitic acid for 8 h. Incorporated C16:0-azide was detected by click chemistry with alkyne-mPEG 5K (200 μM), followed by SDS–PAGE and WB for GPX4. Results are representative of three independent experiments. **f–h**. same as in a, for **f,** HEK WT (Ctrl) or ZDHHC20-KO cells, recomplemented with empty vector or short/long ZDHHC20 or **h,** U2OS cells left untreated or treated for 4 h with 50 μM H₂O₂. For f, Myc-ZDHHC20 expression was confirmed by WB. The levels of ^3^H-palm incorporation were set to 100% in Control-WT cells for g, or untreated control cells, in h. Results are mean ± SEM, and each dot represents an independent experiment (g, n=4 and h, n=6). *p* values comparing to Ctrl or as indicated by, g, two-way ANOVA, Tukey’s test, and h, Student’s t-test. **i.** Cellular glutathione measured in U2OS cells, siCtrl or siZDHHC20 (72 h) alone or recomplemented with short/long ZDHHC20 (24 h). Data are mean ± SEM from one of three independent experiments (n=6). *p* values by, two-way ANOVA, Tukey’s test **j.** Viability assessed using Promega Glo ATB detection in U2OS cells as in i, treated as in h. Data are mean ± SEM from one of three independent experiments (n=36 replicates). *p* values by, two-way ANOVA, Tukey’s test **k.** Proposed Model: Under homeostasis, ATF2–SETDB1–KAP1–ZNF354A repress antioxidant genes. Lipid peroxide accumulation activates p38/JNK signaling, leading to ZNF354A dissociation and induction of antioxidant responses.

We next tested the ability of the long isoform ZDHHC20L to modify GPX4 and found that it was more efficient than the canonical enzyme in promoting GPX4 acylation (Fig. 8fg). Given the stress-induced expression of ZDHHC20L and its proposed role in redox homeostasis, we next asked whether oxidative stress conditions could modulate GPX4 acylation. Remarkably, H₂O₂ treatment had a major effect on GPX4 acylation, as evidenced by a marked increase in ³H-palmitate incorporation into GPX4 (Fig. 8h).

Because GPX4 activity depends on GSH, and its knockdown increases GSH levels (Fig. 7h), we asked whether ZDHHC20 influences cellular GSH levels. Similar to GPX4 and ZNF354A, ZDHHC20 knockdown elevated intracellular GSH (Fig. 8i). This redox imbalance was fully corrected by re-expression of ZDHHC20L, whereas the canonical isoform had a more modest effect (Fig. 8i). Importantly, ZDHHC20L conferred a marked survival advantage: loss of ZDHHC20 impaired cell viability, consistent with recent findings^86^, which was rescued by ZDHHC20L and only partially by the canonical form (Fig. 8j). Finally, ZDHHC20L-expressing cells were significantly more resistant to H₂O₂-induced oxidative stress when compared to cells expressing the canonical isoform or lacking ZDHHC20 (Fig. 8j).

Altogether, these findings demonstrate that ZDHHC20 contributes to cellular redox homeostasis, and that ZDHHC20L outperforms the canonical isoform. This identifies ZDHHC20L as a functional effector of the LORD pathway that protects cells against oxidative stress.

## Discussion

This study uncovers an epigenetically controlled stress-response pathway that protects cells from the harmful accumulation of membrane lipid peroxides. We term this pathway LORD (Lipid Oxygen Radical Defense) (Fig. 8k). In addition to regulating the expression of antioxidant genes, LORD also regulates the activity—but not the transcription—of GPX4, enhancing the cell’s ability to detoxify lipid peroxides, preserve membrane integrity, and modulate susceptibility to ferroptosis.

Under basal conditions, the transcriptional regulator ZNF354A, a central LORD component, represses numerous antioxidant defense genes, acting as a controller of lipid quality and membrane damage repair. ZNF354A is ubiquitously expressed, as shown by GTEx Portal data (https://www.gtexportal.org/home/), with its lowest expression in the liver—a tissue naturally exposed to oxidative stress, potentially reflecting a reduced need to repress antioxidant programs. In contrast, expression is high in the kidney, consistent with early reports in rats, where it was identified as Kid-1 (kidney, ischemia, and developmentally regulated gene 1)^89–91^. Interestingly, in the rat kidney, ZNF354A expression drops to undetectable levels upon reperfusion after ischemia, suggesting LORD pathway activation during reoxygenation. Our genomic liftover analysis indicates that the mouse ortholog ZFP354A may also bind near antioxidant response genes, making it important to examine whether these regulatory networks are conserved across species and shaped by the distribution of transposable elements, in particular LINE-1.

Mechanistically, ZNF354A is part of a repressive complex that includes the global epigenetic regulator KAP1and the H3K9 methyltransferase SETDB1, as for many KZFPs. A crucial non-canonical component of this complex is the transcriptional activator ATF2, which plays a central role in relieving repression in response to stress. Upon lipid peroxide accumulation, ATF2, KAP1 and ZNF354A undergo p38– and JNK-dependent phosphorylation. This triple phosphorylation requirement is intriguing and suggests that LORD is a finely tuned, tightly regulated stress-response pathway—warranting further investigation.

Phosphorylation causes ZNF354A to dissociate from the complex and to be released from chromatin, enabling the expression of antioxidant genes. These genes protect from ferroptosis and possibly other forms of cell death. Notably, not all antioxidant genes fall under LORD transcriptional control. *GPX4* does not. Instead, its activity is modulated by S-acylation through ZDHHC20L and by ZNF354A-dependent control of GSH availability, both downstream of LORD. Thus, although GPX4 gene expression lies outside the LORD network, its enzymatic function is shaped by it. Future studies should, in particular, determine whether selenoprotein synthesis of GPX4 is also under LORD control.

Our findings reinforce the emerging view that KZFPs orchestrate gene regulatory networks across diverse biological processes^35,36,59,92–96^. They also highlight the function of additional factors, such as ATF2, that can dynamically modulate KAP1–KZFP repression complexes in a pathway-specific manner. How ATF2 is selectively recruited to specific KZFP–KAP1 complexes remains an open and important question.

The expression of ZNF354A-repressed genes involves transcriptional activators. For ZDHHC20L, we identified FOXA1, SP1, and NRF2, the latter being a well-established redox regulator. These findings suggest that LORD operates within a broader transcriptional framework. Future work will define the full set of transcription factors involved in the LORD response, detail the dynamics of activation, and test whether repression serves to delay transcriptional responses—allowing coordinated defense to return to homeostasis^2,14^.

The discovery of the LORD pathway addresses a long-standing gap in lipid oxidative stress biology and raises exciting new questions, foremost among them how excessive lipid peroxidation is sensed and how this triggers MAP kinase activation. Our findings also highlight testable therapeutic opportunities, such as increasing ferroptosis sensitivity to eliminate cancer cells or activate the LORD pathway to reduce neurodegeneration and inflammation.

## Acknowledgments

We warmly thank Dr. Martina Begnis, Dr. Julien Duc, Dr. Priscilla Turelli and Prof. Giovanni D’Angelo for helpful discussions throughout the study and comments on the manuscript. This work is supported by a generous foundation represented by CARIGEST SA, and Swiss National Science Foundation grant 310030_192608 to F.G. v.d.G. and grants from the European Research Council (KRABnKAP, No. 268721; Transpos-X, No. 694658), the Swiss National Science Foundation (310030_152879 and 310030B_173337), and the Aclon Foundation to D.T.

## Author contributions

Conceptualization, F.S.M., L.A., R.F., D.T., and F.G.v.d.G.; investigation, F.S.M., L.A., R.F., B.K., C.R., L.B., F.M., D.M., E.P., and O.R.; funding acquisition, F.G.v.d.G.; writing – original draft, F.S.M., L.A., R.F., D.T., and F.G.v.d.G.; writing – review & editing, F.S.M., L.A., R.F., L.B., F.M., D.M., O.R., D.T., and F.G.v.d.G.; resources, B.K. and C.R.

## Declaration of generative AI and AI-assisted technologies in the writing process

During the preparation of this work, the author(s) used ChatGPT in order to improve the language.

## Methods

### Resource and Data availability

All resources and materials will be made available upon reasonable request to Gisou van der Goot. All data supporting the findings of this study are available within the paper, Supplementary Information, Source data and DOI: 10.17632/54dztcj7km.1. All data regarding the human *ZDHHC20* locus was obtained from the Genome browser on Human (GRCh37/hg19) (version hg19); Transcript data obtained from GENCODE Genes track (V4 0lift37) https://genome.ucsc.edu/cgi-bin/hgTrackUi?g=knownGene; Transcription factor binding sites and Chromatin State segmentation obtained from ENCODE https://genome.ucsc.edu/cgi-bin/hgTrackUi?db=hg18&g=wgEncodeBroadHmm and https://genome.ucsc.edu/cgi-bin/hgTrackUi?hgsid=2468617405_fC2FHA661Zobf2VQ1vQVBSTt1M4b&c=chr5&g=wgEncodeReg. Available KZFP ChIP-seq data was obtained from https://tronoapps.epfl.ch/web/krabopedia/

### Cell lines

U2OS (ATCC: HTB_96), MDA-MB-231 (ATCC: HTB_26), K-562 (ATCC: CCL_243), SUDHL4 (ATCC: CRL_2957), SUDHL5 (ATCC: CRL_2958), SUDHL6 (ATCC: CRL_2959), MCF7 (ATCC: HTB_22), SW480 (ATCC: CCL_228), LS1034 (ATCC: CRL_2158), HCT116 (ATCC: CRL_3502), HAP-1 (Horizon Discovery: HZGGHC9001), HAP-1 KO_KAP1 (Horizon Discovery: HZGHC000293c003), HEK293T (ATCC: CRL-3216), HEPG2 (ATCC: HB_8065), Calu-3 (ATCC: HTB_55), 3T3 (ATCC: CRL_1658), VERO-E6 (ATCC: CRL_1586) cells were grown in Dulbecco’s Modified Eagle Medium supplemented with 10% FBS, penicillin, and streptomycin. HELA (ATCC: CCL_2) cells were grown in MEM Media supplemented with 10% fetal calf serum, 1% penicillin–streptomycin, 1% NEAA and 1% glutamine (all media from ThermoFisher Scientific). ZDHHC20 KO HEK293T cells were generated using CRISPR technology ^38^. HEK293T cell lines stably expressing ZNF354A or GFP were generated as previously described^35^.

### Antibodies and Reagents

The antibodies and reagents were acquired from: ATF2 (Abcam: ab_131484; RRID: AB_11156678, rabbit: 1:2000x), ATF2 (Abcam: E243 ab32160, RRID: AB_2243137, rabbit: 1:2000x), phosphor-ATF2 (T69) (Abcam: 131106; RRID: AB_11157608;rabbit:1:1000x), GAPDH (Thermofisher: 398600; RRID: AB_2533438; mouse: 1:4000x), ZDHHC20 (Sigma: SAB4501054; RRID: AB_10744838; rabbit: 1:2000x), HA-HRP (Roche: 112013819001; RRID: AB_390917; rat: 1:5000x), KAP1 (Abcam: ab_109289; RRID: AB_10863057, rabbit: 1:1000x), phosphor-KAP1 S473 was generated in D. TRONO laboratory using the following peptide krsrSgegevsgl (rabbit: 1:500x), Nucleocapside N SARS-CoV-2 (Genetex: GTX135357; RRID: AB_2868464; rabbit 1:4000x), FLAG (Sigma: F3165; RRID: AB_259529; mouse: 1:2000x), phosphor-Threonine-Serine (BD: 612548; RRID: AB_399843; rabbit: 1:1000x), GPX4 for WB (Cell Signalling: 52455; RRID: AB_2924984, rabbit: 1:1000x), GPX4 for IP (MA5-49216, RRID: AB_3074874, mouse 1:2000x), SETDB1 (Novus Biologicals: NBP1_51677; RRID: AB_11002952, mouse: 1:2000x), NRF2 (Abnova: H00004780; RRID:566020, mouse: 1:2000x), V5-HRP (Sigma: V2260; RRID: AB_261857, mouse: 1:2000x), Actin (Millipore: MAB1501; RRID: AB_2223041, mouse: 1:4000x), MAT1A (Abcam: 129176; RRID: AB_11145300, rabbit: 1:2000x), ZNF354A (Santa Cruz: 81140; RRID: AB_1131557, mouse: 1:1000x), MAT1A (Abcam: ab1129176, RRID: AB_11145300, rabbit: 1:1000x), Myc (Millipore: 9E10, 05-419, RRID: AB_309725, mouse: 1:3000x), and HRP-conjugated secondary antibodies (GE-Healthcare).

RSL-3 (Sigma: S8155), Hydrogen Peroxide 30% (Fisher: H/1750/15), Rotenone (Cayman Chemical: CAY-13995), Ferrostatin-1 (Sigma: SML0583), Bodipy 581/591 C11 (Sigma: D3861), Liperfluo (Fisher: L248), Propidium Iodide (Thermofisher: P3566), Cell Viability assay (Promega: G9711), JQ1 (Sigma: SML-1524), Mithramycin (LKT: M3476), Erastin (Millipore: 329600), all resuspended in DMSO and N-acetylcysteine (NAC) (Selleckchem: S1623) resuspended in water, with pH adjusted to 7.4).

### siRNA

For human ZNF354A, 3 different siRNA in 3’-UTR were tested and validated, from Qiagen: ZNF354A (TACATCCTTGAGAGAGATGTA), from Santa Cruz (sc-91974), ZNF354A (GTCACGTGATCATCAGAAA), and from Origene, ZNF354A (GAGGTTTATCCATGGAATTAAATACCT).

siRNAs against all human ZDHHC genes were the same as previously published ^97^. The other following human siRNA with the corresponding target sequences were purchased from Qiagen: ZNF354B (ATCAGTCTTCATTAAGCAACA; CACATTGAAGAGGACTCCTTA; TACATGCTTGAGTGATTTA), ZNF317 (CATGACGGAAATCACACTAAA), ZNF 308 (CAGTTCATTGGCAACCTCATA), ZNF677 (GAGATTGGTTACAGTACAAGA), ZNF793 (ACCCTGGCATGTAAACGTTTA), SETDB1 (TCGGGTGGTCGCCAAATACAA), KAP1 (AGCGTCCTGGCACTAACTCAA), ATF2 (CCGAGGTAGTCCACATACAGAA), NRF2/NFE2L2 (CCCATTGATCTTTCTGATCTA), GPX4 (CGGCTGCGTGGTGAAGCGCTA). The following human siRNA were purchased from Santa Cruz: SP1 (sc-44221), FOXA1 (sc-37930). Mouse siRNA were purchased from Qiagen for mouse ZFP354A (TACCACTTTAATAACATTCAA). As a control siRNA, a sequence targeting the viral glycoprotein VSV-G (5’-ATTGAACAAACGAAACAAGGA-3’) was used. Transfections of 50 nM of siRNA were carried out using *TRANS*IT-X2 (MIRUS), and the cells were analyzed at least 72 h after transfection.

### Western Blotting

Cells were washed three times in 1x PBS at 4°C and lysed in Buffer (1x PBS, 0.5% NP40 and protease inhibitor cocktail; Roche) for 30 min on ice. Lysates were then spin down at 10 000 g, and the protein content of supernatant were determined, the samples were boiled in Laemmli buffer for 5 min before separation via SDS-PAGE and western blotting against the different anti-bodies used in this study. For the analysis of phosphorylated proteins, SuperSep™ Phos-tag™ gels (cat. no. 199-18011) were used. Western blots were developed using the ECL protocol and imaged on a Fusion Solo from Vilber Lourmat. Densitometric analysis was performed using the software Bio-1D from the manufacturer.

### Plasmids

Plasmids expressing HA-tagged ZNF354A-HA, ZNF317-HA, ZNF677-HA, ZNF793-HA, KAP1-HA were obtained from the D. Trono lab ^35^. Plasmids expressing FLAG-ATF2 (RC2118983) were from Origene, myc-SETDB1 (RC2266620) and plasmids expressing SP1-FLAG (25543) and FOXA1-V5 (153109) from Addgene. Mutations to change KAP1-HA to KAP1-FLAG, ZNF354A-HA to ZNF354A-FLAG, FLAG-ATF2 to HA-ATF2, HA-ATF2 6A (T52A-T69A-T71A-T73A-S490A-S498A), HA-ATF2 6D (T52D-T69D-T71D-T73D-S490D-S498D), ZNF354A-HA (S169A), ZNF354A-HA (S169D), ZNF354A-HA (T189A), ZNF354A (T154A), ZNF354A-HA RR (D18R-E27R), ZNF354A RA (E45R-N46A), ZNF354A RR-RA (D18R-E27R-E45R-N46A), KAP1-HA (S473A), KAP1-HA (S473D) and MYC-SETDB1 inactive form (H1224A-C1226A-C1279A-C1280A-C1281A) were generated following Quickchange mutagenesis kit (Agilent) instructions. We generated a pIRES-based expression construct encoding the wild-type cytoplasmic isoform 2 of human GPX4, tagged with Myc and Flag epitopes. This construct includes a UGA codon at position 46, corresponding to the incorporation site of selenocysteine (Sec, U46), as well as the selenocysteine insertion sequence (SECIS) element in the 3′ untranslated region (3′UTR). The SECIS element is a conserved RNA stem-loop structure required for the recoding of the UGA stop codon as selenocysteine during translation, enabling the proper synthesis of selenoproteins.

### Q-RT-PCR

For the real-time PCR, RNA was extracted from a six-well dish using the RNeasy kit (Qiagen). In all, 0.5-1 mg of the total RNA extracted was used for the reverse transcription using random hexamers and superscript II (Invitrogen). A 1:40 dilution of the cDNA was used to perform the real-time PCR using using Applied Biosystems SYBR Green Master Mix on 7900 HT Fast QPCR System (Applied Biosystems) with SDS 2.4 Software. All data (always in triplicate) were normalized to Ct values from three housekeeping (HK) genes ALAS-1, Guss and TBP. Results were expressed as 2^(-ΔΔCt)*100%.

### Oligonucleotides for Quantitative RT-PCR

All QPCR primers, used in this study are described in detail in table below (Supplementary Table 1).

**Supplementary Table 1.**
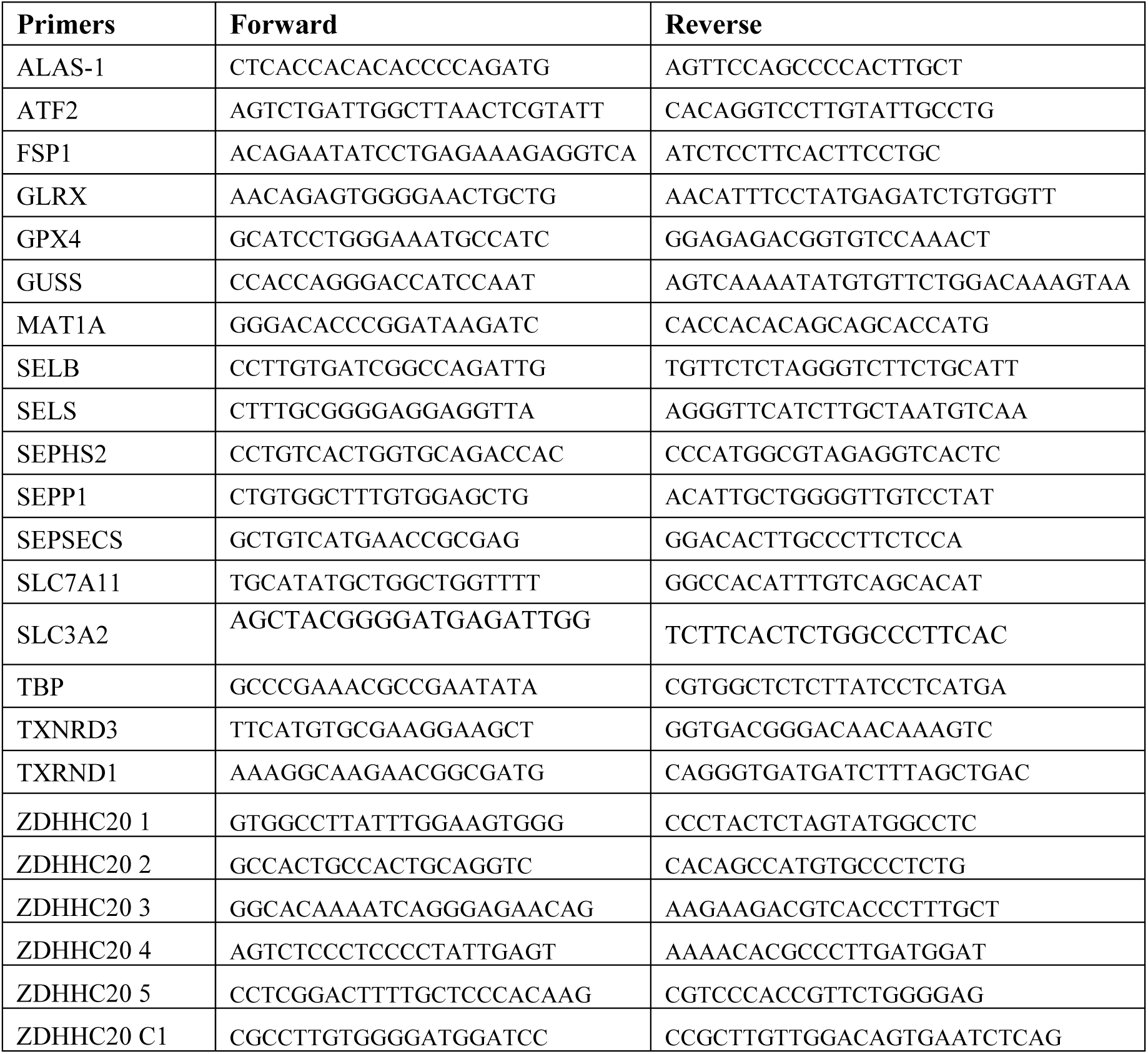
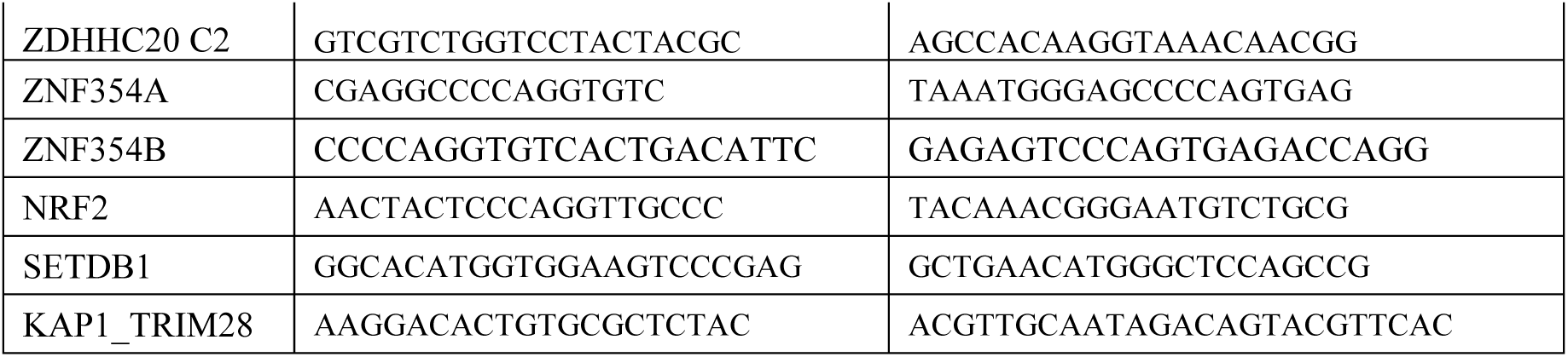
Primers against human genes used in this study.

### Aerolysin purification and cell intoxication

Proaerolysin toxin was produced and purified by our laboratory in Aeromonas salmonicida as previously described (ref). Cells were treated one hour in complete medium with 10 ng/ml of proaerolysin at 37 °C. Cells were washed twice in complete medium and further incubated at 37 °C for indicated times.

### GREAT – analysis

ChIP-seq data for ZNF354A-HA binding sites were obtained from KRABopedia ^58^ and submitted to the Genomic Regions Enrichment of Annotations Tool (GREAT – http://great.stanford.edu/public/html/) ^71^. The parameters for gene regulatory domains for each gene were defined as domain that extends in both directions of the TSS until the nearest TSS up to a maximum extension of 300 kb.

### Human MucilAir assays

MucilAir-human upper respiratory tissues (Epithelix, Geneve, Switzerland) were maintained according to the manufacturer’s protocol. For infection, tissues were washed apically with 200 μl of DPBS, calcium, magnesium (#14040091—Gibco) for 20 min at 37 °C and the basal medium replaced with fresh mucilair medium. Tissues were infected at the MOI of 0.1 PFU (assuming the manufacturer’s estimations of 500000 cells per tissue). Tissues were inoculated with 200 μl of SARS-CoV-2 (passage 3) for 3 h (apically) at 33 °C. The apical inoculum was removed and infected tissues were maintained until harvest. For harvest tissues were lysed using 300 μl lysis buffer for 30 min and processed for western blot.

### Viral Stock production and titration with plaque-based assays

All viral stocks were produced and isolated from supernatants of Vero E6 cells, cultured in T75 culture flasks to a confluency of 80-90%, and infected with an original passage 2 (P2) SARS-CoV-2 virus, for 48 or 72 h, at MOI≈0.05, in 10 ml DMEM supplemented with 2.5% FCS. Original stocks were obtained from the following strain: hCoV-19/Switzerland/GE9586/2020|EPI_ISL_414022|2020-02-27. Passage 3 supernatants were harvested, clear of cell debris by centrifugation (500 *g* 10 min) and filtration (0.45 µm), aliquoted and stored at –80 C. Viral titers were quantified by determining the number of individual plaque forming units after 48 h of infection in confluent Vero E6 cells as described in ^37^.

### SARS-CoV-2 infections

All infections for experimental analysis were done using passage 3 SARS-CoV-2 stocks. Vero E6 cells or CALU-3 cells seeded to a confluency of 90 to 100%, were, washed twice in warm serum-free medium and inoculated with the indicated MOI of SARS-CoV-2, diluted in serum free medium. 1 hour after inoculation cells were washed with complete medium and infection allowed to proceed for the indicated time points in DMEM supplemented with 2.5% FCS, penicillin and streptomycin.

### Drug and toxin treatments

All drug treatments were done for the indicated times in complete culture and/or infection medium. Equivalent volume of Dimethyl sulfoxide (DMSO), used as a solvent, was added for control samples. Specifically, culture media was replaced, cells were washed once in warm culture media and drugs resuspended as indicated were used as: RSL3 (in DMSO – 3 μM, increasing concentrations 0.6 to 10 μM), Ferrostatin-1 (DMSO – 1 – 15 μM), H₂O₂ (0.5 – 100 μM), Rotenone (1 – 10 μM).

### Data mining analysis

UCSC genome browser at ZDHHC20 locus was obtained from (Human version hg19). Specific UCSC genome browser tracks were selected for display in Fig. 1 and 2. Transcript annotation corresponds to GENCODE Genes track (version V4 0lift37); Histone Modifications by ChIP-seq from ENCODE/Stanford/Yale/USC/Harvard display ChIP-seq signals for H3K4me1, H3K4me3, H3K27Ac marks for HepG2 and H3K9me3 for U2OS. Transcription Factor CHIP-seq Peaks track shows transcription factor SETDB1, and KAP1binding sites for U2OS and K562 cells based on ChIP-seq experiments from ENCODE. KZFP CHIP-seq data show binding sites for different KZFP proteins obtained from KRABopedia (https://tronoapps.epfl.ch/web/krabopedia/) (https://doi.org/10.1101/2023.02.27.530095)

### Chromatin Immunoprecipitation (ChIP) followed by qPCR or sequencing (ChIP-seq)

K562 cells (30 million) were cross-linked at room temperature for 10 minutes by adding formaldehyde to a final concentration of 1%, followed by quenching with glycine Tris 200mM. Cells were washed twice with PBS, pelleted, and stored at –80°C. Pellets were sequentially lysed in LB1 (50 mM HEPES-KOH pH 7.4, 140 mM NaCl, 1 mM EDTA, 0.5 mM EGTA, 10% glycerol, 0.5% NP-40, 0.25% Triton X-100, protease inhibitors), LB2 (10 mM Tris-HCl pH 8.0, 200 mM NaCl, 1 mM EDTA, 0.5 mM EGTA, protease inhibitors), and SDS shearing buffer (10 mM Tris-HCl pH 8.0, 1 mM EDTA, SDS 0.15%, protease inhibitors). Chromatin was sonicated (Covaris: 5% duty, 200 cycles, 140 PIP, 20 min 25min) to obtain DNA fragments of 100-300bp. ChIP was performed anti-HA magnetic beads (Pierce: 88837). Chromatin was adjusted for immunoprecipitation using 1% Triton X-100 and 150mM NaCl then transferred on beads, followed by overnight incubation at 4°C. Immunoprecipitated complexes were washed sequentially with Low Salt Wash Buffer (10 mM Tris-HCl pH 8.0, 1 mM EDTA, 150 mM NaCl, 1% Triton X-100, 0.15% SDS, protease inhibitors), High Salt Wash Buffer (10 mM Tris-HCl pH 8.0, 1 mM EDTA, 500 mM NaCl, Triton X-100, 0.15% SDS, protease inhibitors), LiCl Buffer (10 mM Tris-HCl pH 8.0, 1 mM EDTA, 0.5 mM EGTA, 250 mM LiCl, 1% NP-40, 1% NaDOC, protease inhibitors), and TE buffer. DNA was eluted and purified using QIAGEN columns. Experiments were performed in biological duplicates, Samples were send for sequencing and ChIP-QPCR experiments were independently repeated 3 or 4 times for each depicted primer. ChIP and input DNA were analyzed by qPCR using specific primers and sequencing. Enrichment was calculated as a percentage of input or relative to a control region using the ΔΔCt method. Primers used for QPCR are listed below:

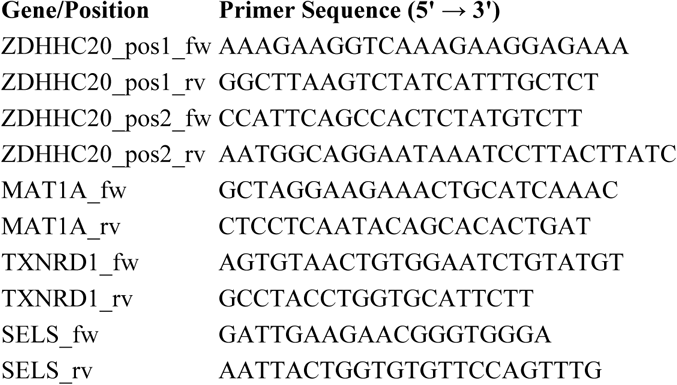

### Library preparation and sequencing

DNA concentrations were determined using a Qubit fluorometer (Thermo Fisher Scientific) following the manufacturer’s protocol. Chromatin immunoprecipitated DNA was processed using the NEBNext Ultra II DNA Library Prep Kit (New England Biolabs, Version 7.0_9/22) according to the manufacturer’s instructions (internal experiment code GFX016). Input DNA amounts varied between 1.5 ng and 5 ng, as specified in the corresponding sample table. Given the absence of initial DNA profile data, a positive control was included during library preparation. Library concentrations were measured using Qubit High Sensitivity (HS) DNA quantification, and fragment size distributions were assessed using a TapeStation TS4200 (Agilent Technologies). All libraries exhibited expected fragment profiles and sufficient yield for sequencing. Prepared libraries were sequenced on the Element Biosciences Aviti platform in a paired-end 150 bp (PE150) run (run ID: AVT0149). Sample-specific identifiers and sequencing metadata are detailed in the supplementary data. Adapter sequences were trimmed using Cutadapt vX.X with TruSeq adapter sequences:

Read 1:AGATCGGAAGAGCACACGTCTGAACTCCAGTCA;

Read2: AGATCGGAAGAGCGTCGTGTAGGGAAAGAGTGT

### ChIPseq analysis

To identify and merge significant peaks across multiple conditions, we utilized BEDTools^98^ (v2.30.0). Initially, overlapping peaks from multiple BED files were merged using the bedtools multiinter function. To ensure a more comprehensive peak detection, an alternative strategy was applied, merging all available BED files in the directory. Following peak merging, signal intensity filtering was performed using deepTools (v3.5.1). The computeMatrix function extracted signal values from bigWig (BW) files over predefined peak regions (Filt_GFP.bed). A threshold of 15 was applied to exclude peaks with excessively high signal intensities. To visualize the signal distribution, a heatmap was generated using plotHeatmap, employing the Oranges_r colormap. Based on the filtered set (sorted_regionsFilt12.bed), another round of peak merging was conducted to ensure that only the most relevant peaks were retained. This final merged peak list contained only peaks present in HEK293T and those retained in K562. To further refine peak selection and exclude aberrant values, we applied an additional filtering step with a stricter intensity threshold (--maxThreshold 150). A final heatmap was generated to visualize the refined peak set. A final heatmap was generated to visualize the refined peak set. Ultimately, the final set of high-confidence peaks used for downstream analysis was stored in sorted_regions_3.bed, representing the most relevant and biologically meaningful peak regions across conditions.

### RNAseq

RNA quality was controlled on the TapeStation 4200 (Agilent), which confirmed their integrity. Libraries for mRNA-seq were prepared with the Stranded mRNA Ligation method (Illumina), according to manufacturer’s instructions, starting from 800ng RNA. Libraries, all bearing unique dual indexes, were subsequently loaded at 100 pM on a Novaseq 6000 flow cell (Illumina) and sequenced according to manufacturer instructions, yielding pairs of 60 nucleotides reads at a depth of about 70 mio reads pairs per sample. Reads were trimmed of their adapters with bclconvert v00.000.000.3.9.3 (Illumina) and quality-controlled with fastQC v0.11.9. FastQ Screen v0.14.0 tool was used for screening FASTQ files reads against multiple reference genomes.

### RNAseq processing and analysis

Sequencing reads were mapped to the human hg19 genome using hisat2 [D. Kim, B. Langmead, and S. L. Salzberg, “HISAT: a fast spliced aligner with low memory requirements,” *Nature Methods*, vol. 12, no. 4, pp. 357–360, Mar. 2015.] with parameters: hisat2 –k 5 –-seed 42 –-rna-strandness RF. Counts on genes were generated using featureCounts [Y. Liao, G. K. Smyth, and W. Shi, “featureCounts: an efficient general purpose program for assigning sequence reads to genomic features,” *Bioinformatics*, vol. 30, no. 7, pp. 923–930, Apr. 2014.]. Only uniquely mapped reads were used for counting on genes. Normalization for sequencing depth was done using the TMM method as implemented in the limma package of Bioconductor [Gentleman et al., 2004]. Differential gene expression analysis was performed using voom (Law et al., 2014) as it has been implemented in the limma package of Bioconductor. A gene was considered to be differentially expressed when the fold change between groups was bigger than 2 and the p-value was smaller than 0.05. A moderated t-test (as implemented in the limma package of R) was used to test significance. P-values were corrected for multiple testing using the Benjamini-Hochberg’s method^97^. Gene set enrichment analysis (GSEA) was performed using the ClusterProfiler package in R. The analysis utilized a ranked list of genes and gene set collections downloaded from MsigDB. The GSEA was conducted with the following parameters: the number of permutations was set to 10,000, minimum gene set size was 10, maximum gene set size was 800.

### Flow cytometry analysis of membrane permeability and lipid peroxidation

U2OS cells (1 × 10^6 per well, 6-well plate) were transfected with siRNA for 72 h. For Liperfluo staining (Dojindo), cells were treated with 5 µM RSL-3 for 24 h, washed twice with EBSS-10 µM HEPES, and incubated in 1 mL EBSS-HEPES with 5 µM Liperfluo for 30 min at 37 °C. Cells were washed, trypsinized, washed in 1% FBS EBSS-HEPES to remove trypsin, resuspended in 500 µL EBSS-HEPES, filtered, and stained with propidium iodide (1 µg/mL) prior to flow cytometry. For C11-BODIPY581/591 (Invitrogen) staining, siRNA-treated cells were washed twice in HBSS and incubated in 1 mL HBSS with 10 µM probe for 30 min at 37°C. Excess dye was removed and cells were treated with 5 µM RSL-3 for 4 h in complete medium. Cells were then processed as above but stained with DAPI (1 µg/mL). Flow cytometry was performed on an LSRII or LSR Fortessa (BD Biosciences). Oxidized Liperfluo and BODIPY were detected using 488 nm excitation and 530/30 nm emission; PI and reduced BODIPY were detected using 561 nm excitation and 610/20 nm emission. At least 10,000 live events (PI/DAPI-negative) were acquired per sample and data were analyzed using FlowJo.

### CLICK-PEG Assay

The protocol was adapted from ^88^. Twenty-four hours after transfection, cells were washed with serum-free medium containing 1 mg/mL BSA and incubated for various times with either 100 µM unlabeled palmitic acid (control) or 100 µM palmitic acid-azide. After labeling, cells were washed twice with cold PBS and lysed in 50 mM Tris-HCl (pH 8.0), 0.5% SDS. Lysates were subjected to a click reaction with 5 kDa mPEG-alkyne (JKA-3177, Sigma), CuSO₄, TBTA, and ascorbic acid for 1 hour at room temperature. The reaction was stopped by boiling the samples in 4× SDS–BME sample buffer (SB4×).

### Intracellular glutathione measurements

Intracellular reduced glutathione levels were measured using the QuantiChrom Glutathione Assay Kit (DIGT-250, BioAssay Systems), following the manufacturer’s instructions. Briefly, 1 × 10⁶ cells were collected and processed for the colorimetric detection of reduced glutathione. The assay is based on the reaction of glutathione with 5,5′-dithiobis-(2-nitrobenzoic acid) (DTNB), which yields a yellow-colored product. Absorbance was measured at 412 nm, and values were directly proportional to the glutathione concentration in the sample.

### ^3^H-palmitic acid radiolabeling experiments

HeLa cells were transfected or not with different constructs, incubated for 2 h in Glasgow minimal essential medium (IM; buffered with 10 mM HEPES, pH 7.4) with 200 µCi/ml of ^3^H palmitic acid (9,10-^3^H[N]) (American Radiolabeled Chemicals, Inc.). The cells were washed, and incubated in DMEM complete medium for the indicated time of chase or directly lysed for immunoprecipitation with the indicated antibodies. After immunoprecipitation, washed beads were incubated for 5 min at 90°C in reducing sample buffer prior to 4−20% gradient SDS-PAGE. Gels were incubated 30 min in a fixative solution (25% isopropanol, 65% H2O, 10% acetic acid), followed by a 30 min incubation with the signal enhancer Amplify NAMP100 (GE Healthcare). The radiolabeled products were imaged using a Typhoon phosphoimager and quantified using a Typhoon Imager (ImageQuanTool, GE Healthcare). The images shown for ^3^H-palmitate labeling were obtained using fluorography on film.

### DSS-induced mice colitis

Six wild-type C57BL/6j mice (8-week-old males) were used. Three (n = 3) mice were used as controls and three mice were given 3% dextran sulfate sodium in the drinking water for 7 days, then switched to regular drinking water for 3 days. Three Control 8-week male were given drinking water that did not contain DSS. During the 10-day experiment, mice were weighted and disease activity index (DAI) was performed daily based on stool consistency, presence of occult blood and body weight loss. On day 10, mice were euthanized with injection of pentobarbital and bled via cardiac puncture, and colon and intestines were collected. For animal experimentation, all procedures were performed according to protocols approved by the Veterinary Authorities of the Canton Vaud and according to the Swiss Law (license VD 3497, EPFL).

### Quantification and Statistical Analysis

All experiments were conducted independently a minimum of three times. Unless indicated otherwise, each data point corresponds to a separate experiment yielding consistent outcomes. Statistical evaluations were carried out using Prism software, with graphical representations and detailed statistical information available in the figure legends. For ANOVA, p-values were calculated using post hoc tests: Tukey’s and Sidak’s for multiple comparisons between groups, and Dunnett’s for comparisons against a control condition. RNA-seq and ChIP-seq analyses were performed using distinct bioinformatics pipelines, as outlined in the Materials and Methods section, to ensure appropriate data handling and statistical evaluation specific to each dataset.

## Supplementary information

**Supplementary Table 2.** Gene Set Enrichment Analysis-GSEA dot plot of ZNF354A upregulated enriched GO-terms depicted in Fig 3C.

| Ontology | setSize | pvalue |
| --- | --- | --- |
| GOBP_CHEMICAL_HOMEOSTASIS | 772 | 0.000120555 |
| GOBP_REGULATION_OF_RESPONSE_TO_EXTERNAL_STIMULUS | 713 | 0.000122549 |
| GOCC_VACUOLE | 711 | 0.000122594 |
| GOBP_RESPONSE_TO_LIPID | 669 | 0.000122956 |
| GOBP_DEFENSE_RESPONSE_TO_OTHER_ORGANISM | 704 | 0.000123062 |
| GOBP_RESPONSE_TO_CYTOKINE | 698 | 0.000123183 |
| GOBP_REGULATION_OF_IMMUNE_RESPONSE | 599 | 0.000125423 |
| GOBP_INFLAMMATORY_RESPONSE | 550 | 0.000127146 |
| GOBP_INNATE_IMMUNE_RESPONSE | 550 | 0.000127146 |
| GOBP_CYTOKINE_PRODUCTION | 550 | 0.000127146 |
| GOBP_PROCESS_UTILIZING_AUTOPHAGIC_MECHANISM | 510 | 0.0001287 |
| GOCC_ENDOSOME_MEMBRANE | 479 | 0.000129433 |
| GOBP_REGULATION_OF_DEFENSE_RESPONSE | 464 | 0.000129836 |
| GOCC_VACUOLAR_MEMBRANE | 417 | 0.000131752 |
| GOBP_RESPONSE_TO_BACTERIUM | 422 | 0.000131961 |

**Supplementary Table 3.**
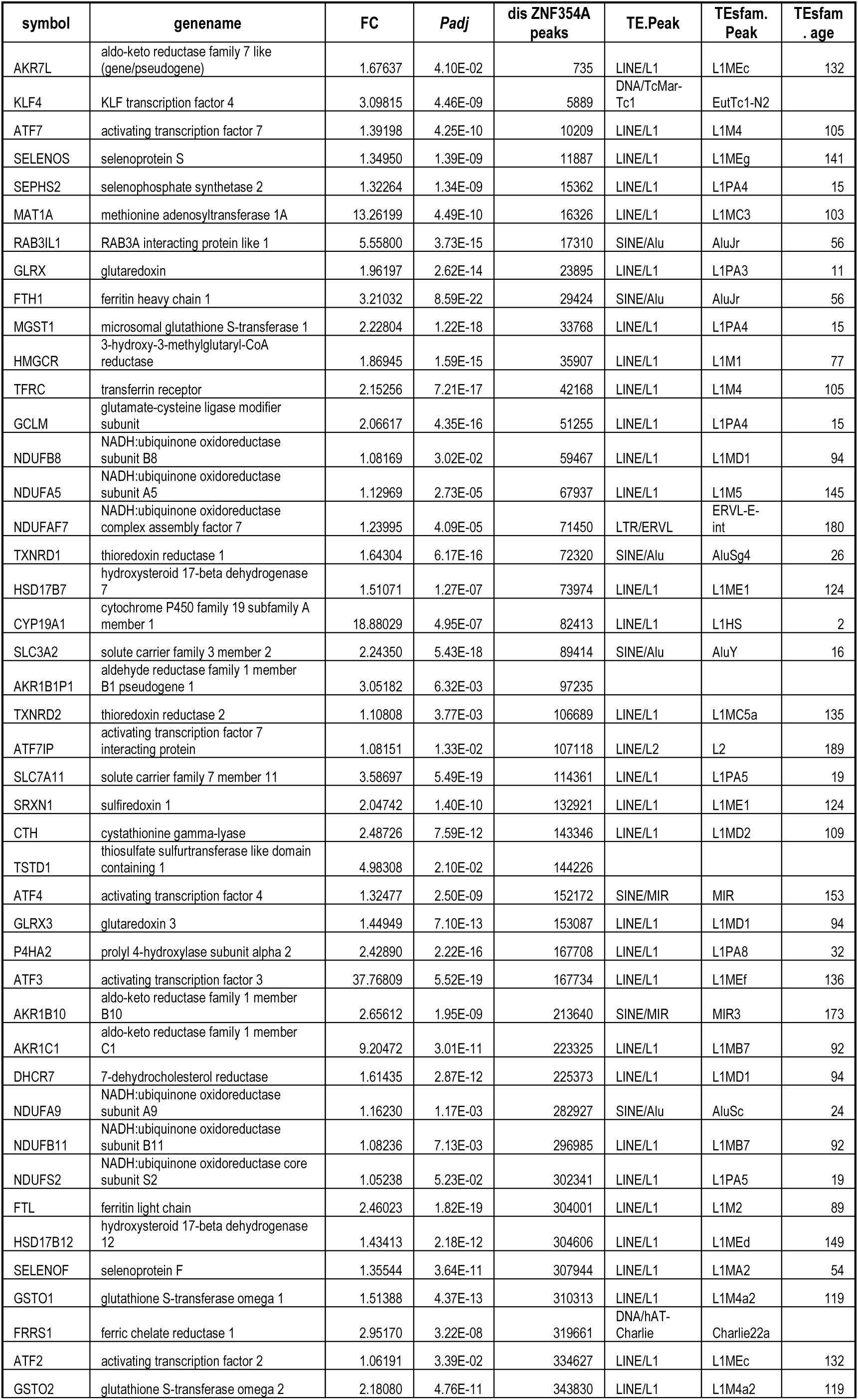
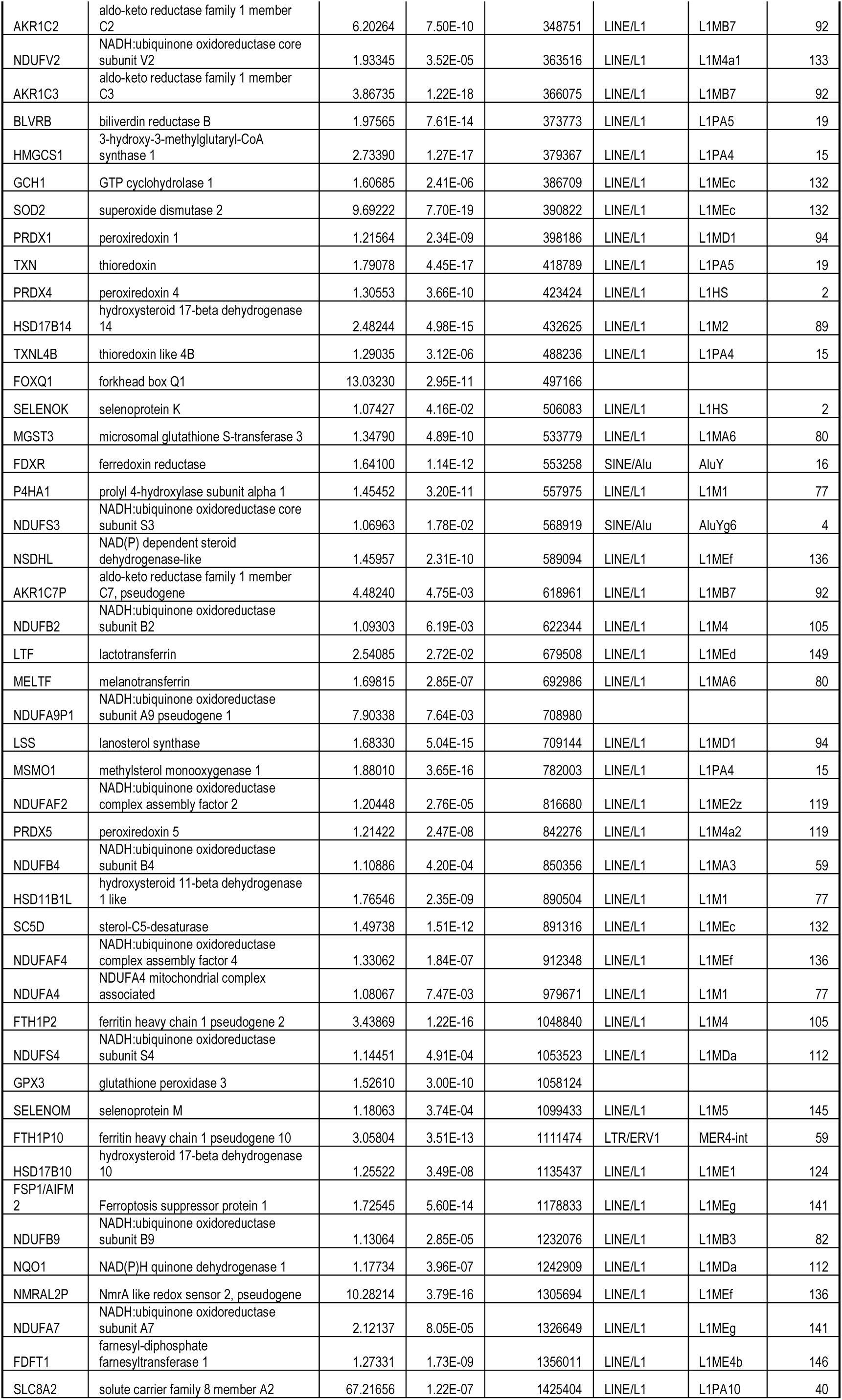
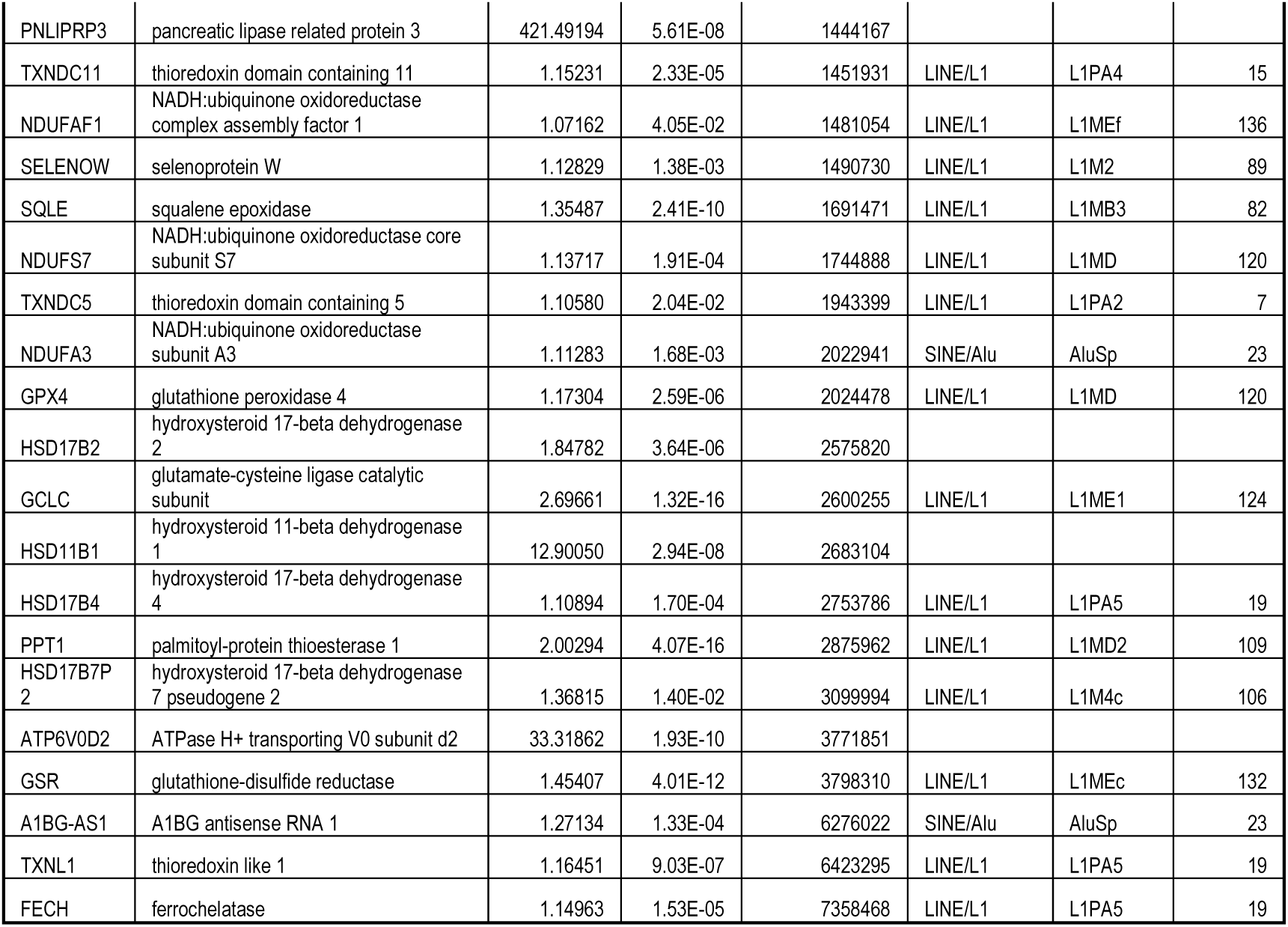
Selected Antioxidant genes up-regulated upon siZNF354A. Fold enrichment, adjusted P values, distance to nearest ZNF354A peak, and corresponding overlapping TE (with Subfam and age) are indicated.

